# Moisture structures litter faunal communities through productivity and trait filtering effects

**DOI:** 10.64898/2026.09.18.752524

**Authors:** N.J. Butterworth, N. Porch, A.D. Barnes, A. Carlesso, M.A. McGeoch, N.P. Murphy, M. Dawson, H. Gibb

## Abstract

Litter invertebrates mediate a substantial proportion of decomposition and nutrient cycling in terrestrial ecosystems, yet how moisture availability structures their communities remains poorly understood. We propose that moisture availability could shape litter faunal communities through two alternate pathways: (1) a productivity effect: greater productivity and detrital inputs in wetter forests should increase litter-faunal abundance, (2) a trait effect: phylogenetic differences in water balance should filter communities according to taxon-specific moisture tolerances. We tested these expectations across 100 forest sites spanning a moisture gradient in south-eastern Australia, from dry sclerophyll forest to cool temperate rainforest. Measuring 11 focal invertebrate taxa, total litter-faunal abundance increased threefold across the moisture gradient, from ∼400,000 individuals per hectare of dry forest to ∼1.2 million individuals per hectare of rainforest – consistent with greater resource availability in wetter and more productive habitats. However, this overall abundance trend masked pronounced variation in community composition across habitat types. While moisture-sensitive crustaceans and myriapods (particularly Isopoda and Chilopoda) were concentrated in rainforest litter, two groups of hexapods (Blattodea and Embioptera) were most abundant in dry forest litter and declined sharply toward rainforest habitat. These contrasting responses indicate that dry forests do not simply represent depauperate versions of wet-forest communities but harbour their own distinct and abundant litter invertebrate assemblages. Thus, moisture structures litter faunal communities via both proposed mechanisms: (1) a productivity effect and (2) a trait filtering effect. Given that moisture fundamentally structures litter faunal communities, shifts in forest moisture regimes under climate change may substantially alter detritivore communities, with consequences for decomposition processes and nutrient cycling globally.

## Introduction

Water availability is a fundamental determinant of forest ecosystem structure and function (Schuur et al. 2001; Malhi et al. 2015; Bastin et al. 2017; Moore et al. 2017; Bennett et al. 2020; Ma et al. 2021; Zhang-Zheng et al. 2024). Across the globe, forests span a broad hydrological spectrum, from the arid Baobab forests of Senegal (≤400mm of annual rainfall) to the saturated Chocó–Darién rainforests of Colombia and Ecuador (>9,000mm of annual rainfall). Across this gradient, wetter forests generally support higher plant biomass (Silvertown et al. 1994; Keith et al. 2009; Pan et al. 2013; Moore et al. 2018; Aponte et al. 2020), elevated rates of litterfall and decomposition (Murphy et al. 1998; Thomas et al. 2014; Fensham et al. 2024), and increased net primary productivity (Kicklighter et al. 1999; Moore et al. 2018; Muller-Landau et al. 2021). Because moisture so strongly regulates the functioning and resilience of forest ecosystems (Allen et al. 2015; Millar & Stephenson 2015; Siedl et al. 2011; Liu et al. 2024) ongoing shifts in precipitation regimes and drought frequency are expected to restructure forest communities and alter ecosystem processes worldwide (Maestre et al. 2015; Ochoa-Hueso 2018; Aguirre-Gutiérrez et al. 2019; Denissen et al. 2022; Müller & Bahn 2022; Moss et al. 2024). To predict the outcomes of these changes, we must understand how the ecological communities that sustain critical ecosystem processes in forests respond across forest moisture gradients.

A critical ecosystem process that is particularly sensitive to moisture is decomposition (Cisneros-Dozal et al. 2007; Powers et al. 2009; Petraglia et al. 2018; Joly et al. 2023). Approximately 90% of net primary productivity enters the detrital pathway, and more than half of this is returned to soils through the decomposition of plant litter (Wardle et al. 2004; Cebrian 1999). This process is mediated by microbial, fungal, and invertebrate decomposers, all of which are highly sensitive to moisture availability (David 2014; Joly et al. 2018; Njoroge et al. 2021; Prescott & Vesterdal 2021; Eisenhauer et al. 2023; Brockett et al. 2012; García-Palacios 2013; Sagi & Hawlena 2023; Veldhuis et al. 2016; Wall et al. 2008; Tan et al. 2020; Torsekar et al. 2024; Luan et al. 2024). Global syntheses estimate that litter invertebrates alone mediate 24-48% of forest litter decomposition (Sagi & Hawlena 2023; Zeng et al. 2024). Their contribution may be particularly important in xeric environments where microbial and fungal activity becomes constrained by low moisture (Cheli et al. 2022; Seely & Luow 1980; Allison et al. 2013; Manzoni et al. 2012; Maestre et al. 2015; Coleine et al. 2024; Silva et al. 1985; Cepeda-Pizarro & Whitford 1989; Njoroge et al. 2021; Sagi & Hawlena 2023; Torsekar et al. 2024). Consequently, as drought and disturbance intensify under climate change, the persistence and functioning of litter invertebrate communities may become increasingly crucial for ensuring that nutrient cycling and forest ecosystem resilience are maintained (Butler et al. 2019; Veldhuis et al. 2016; Sagi & Hawlena 2023; Torsekar et al. 2024; Luan et al. 2024) – particularly in dryer forests where litter detritivores may play an exceedingly important role.

One of the primary ways litter invertebrates should respond to moisture is through changes in population densities (Janzen & Schoener 1968). This is in line with plant biomass patterns, because wetter forests generally produce more litter biomass (Thomas et al. 2014), exhibit faster decomposition rates (Murphy et al. 1998; Cisneros-Dozal et al. 2007; Zhang et al. 2008; Thomas et al. 2014; Fensham et al. 2024), and maintain more stable microclimatic conditions that reduce desiccation stress and maximise activity of soil fauna (Kaspari & Weiser 2000; Collison et al. 2013) – all of which should favour greater densities of litter invertebrates. Importantly, changes in the number of litter invertebrates have direct implications for ecosystem functioning because the abundance and density of litter consumers should strongly influence the rate and energetics of detrital resource consumption, decomposition, and nutrient turnover (Eisenhauer et al. 2023; Allison et al. 2013; Carbone et al. 2007; Hobbs 2024; Klemmer et al. 2012; del Campo et al. 2025). Evidence at the micro-scale suggests that litter microbe abundance does increase with moisture (Allison et al. 2013; Maestre et al. 2015) and for invertebrates this notion is supported by numerous studies showing positive correlations between precipitation or moisture and invertebrate abundance or biomass (Janzen & Schoener 1968; Frith & Frith 1990; Chikoski et al. 2006; Davis et al. 2006; Landesman et al. 2011; Andriuzzi et al. 2020; Martin et al. 2024; Anderson & Smith 2000; Shaftel et al. 2021; Tanaka & Tanaka 1982; González & Seastedt 2000). Thus rainforests, which are characterized by high productivity, stable moisture, and buffered understory climates, should support the highest abundances of litter invertebrate taxa.

However, the responses of soil and litter invertebrates to moisture are not always positive. Taxa can exhibit neutral (Taylor et al. 2004; Lensing et al. 2005) or even negative responses to increasing moisture availability (Levings & Windsor 1985; Chikoski et al. 2006; Andriuzzi et al. 2020; Homet et al. 2021). This variability arises because moisture is not only a resource gradient influencing productivity but is also an environmental filter (*sensu* Kraft et al. 2014) that selects for taxa differing in their tolerance to desiccation or saturation (which can inundate resources, cause drowning, or increase susceptibility to parasites; Grant & Villani 2003; de Groot et al. 2025). The specific moisture requirements of terrestrial invertebrates are partially constrained by higher taxonomic affinities (Little 1983; Chown & Nicolson 2004) (Table 1) and reflect the timing of terrestrial colonization events which occurred under distinct climates and via fundamentally different adaptations to cope with the major new risk of water loss (Hurley 1968; Edney 1968; Schweizer et al. 2019; Warburg 1965) – myriapods and arachnids by the Silurian (416-443 Mya), hexapods in the Devonian (398-416 Mya), gastropods in the Carboniferous (358-298 Mya), and crustaceans (Amphipoda and Isopoda) between the Carboniferous and Cenozoic (300-66 Mya) (Dunlop et al. 2013; Vermeij & Watson-Zink 2022; Myers 2022; Thorpe 2024). Generally, the Crustacea, Onychophora, Gastropoda, and Myriapoda are considered hygrophilic, i.e., amphipods can die within minutes in the absence of moisture (Williamson 1951). In contrast, the Arachnida and Hexapoda are considered more adapted to xeric conditions, i.e., the Namib desert tenebrionids (Hexapoda: Coleoptera) which thrive in the driest environments on Earth (Ahearn 1970; Ward & Seely 1996; Roberts et al. 2025). Thus, rather than only increasing total invertebrate abundances, forest moisture gradients should also environmentally filter xeric-versus mesic-adapted lineages. However, there are many examples that defy the above generalities, including xeric-adapted crustaceans (Warburg 1965), myriapods (Crawford et al. 1987), and gastropods (Schmidt-Nielsen et al. 1971), and hygrophilic hexapods (Kellerman et al. 2012; Slatyer & Schoville 2016) and arachnids (Lapinski & Tschapka 2014). So, it remains unresolved whether litter invertebrate abundance does scale predictably along forest moisture gradients, implying a similar but deprived community in dry forests, or remains relatively constant with compositional variation.

**Table 1.**
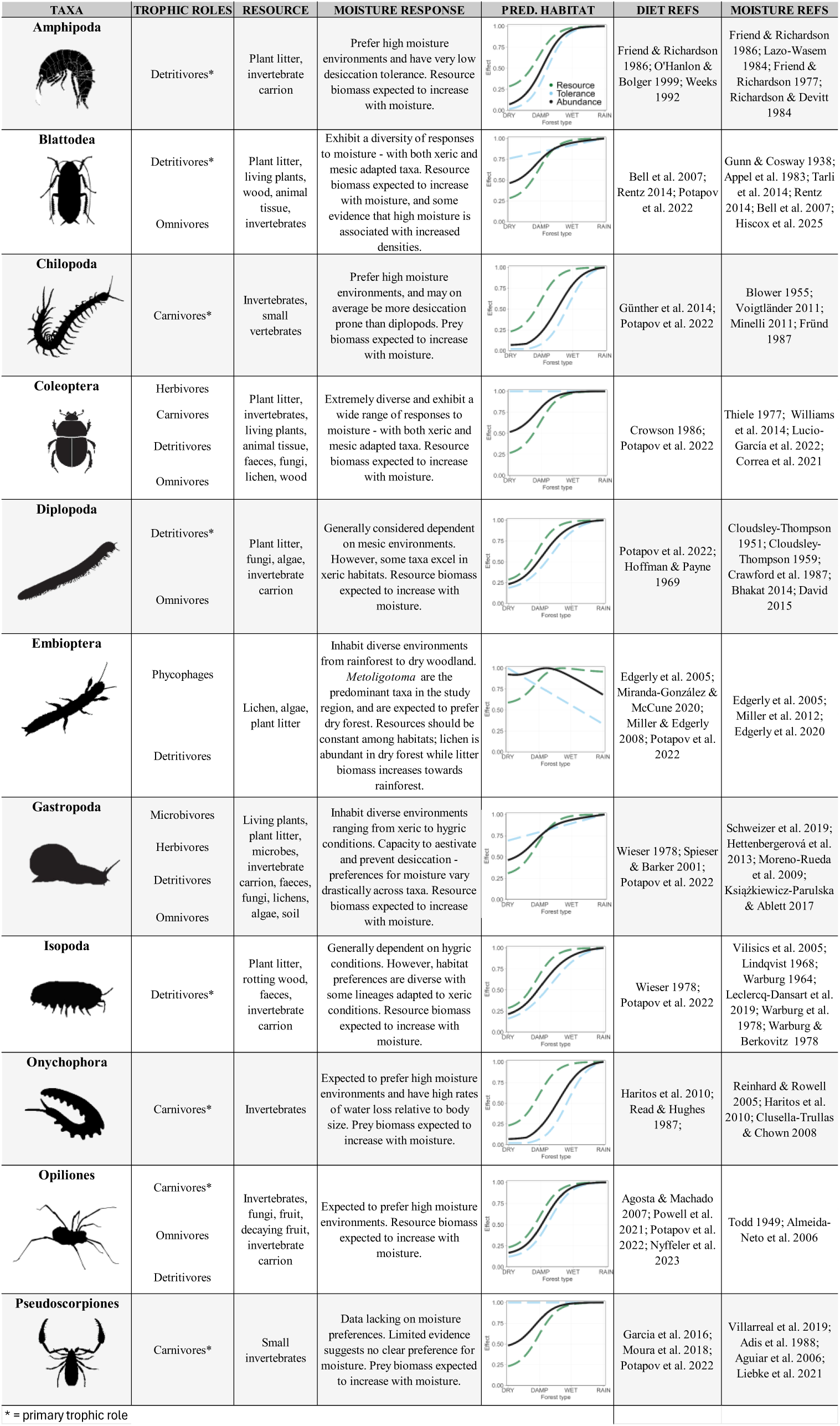
Trophic roles, dietary resources, and expected moisture responses of the 11 focal invertebrate taxa. The moisture response column summarises qualitative expectations based on how each taxon’s moisture tolerance and resource base vary with moisture availability. The predicted habitat column shows, for each taxon, hypothesised resource availability (green dashed line) and moisture tolerance (how physiologically suited the taxon is to the moisture conditions; blue dashed line) across the gradient from dry forest to rainforest, with predicted relative abundance (black solid line) calculated as the square root of the product of resource availability and moisture tolerance.

| TAXA | TROPHIC ROLES | RESOURCE | MOISTURE RESPONSE | PRED. HABITAT | DIET REFS | MOISTURE REFS |
| --- | --- | --- | --- | --- | --- | --- |
| **Amphipoda** | Detritivores* | Plant litter, invertebrate carrion | Prefer high moisture environments and have very low desiccation tolerance. Resource biomass expected to increase with moisture. |  | Friend & Richardson 1986; O'Hanlon & Bolger 1999; Weeks 1992 | Friend & Richardson 1986; Lazo-Wasem 1984; Friend & Richardson 1977; Richardson & Devitt 1984 |
| **Blattodea** | Detritivores*<br>Omnivores | Plant litter, living plants, wood, animal tissue, invertebrates | Exhibit a diversity of responses to moisture - with both xeric and mesic adapted taxa. Resource biomass expected to increase with moisture, and some evidence that high moisture is associated with increased densities. |  | Bell et al. 2007; Rentz 2014; Potapov et al. 2022 | Gunn & Cosway 1938; Appel et al. 1983; Tari et al. 2014; Rentz 2014; Bell et al. 2007; Hiscox et al. 2025 |
| **Chilopoda** | Carnivores* | Invertebrates, small vertebrates | Prefer high moisture environments, and may on average be more desiccation prone than diplopods. Prey biomass expected to increase with moisture. |  | Günther et al. 2014; Potapov et al. 2022 | Blower 1955; Voigtländer 2011; Minelli 2011; Fründ 1987 |
| **Coleoptera** | Herbivores<br>Carnivores<br>Detritivores<br>Omnivores | Plant litter, invertebrates, living plants, animal tissue, faeces, fungi, lichen, wood | Extremely diverse and exhibit a wide range of responses to moisture - with both xeric and mesic adapted taxa. Resource biomass expected to increase with moisture. |  | Crowson 1986; Potapov et al. 2022 | Thiele 1977; Williams et al. 2014; Lucio-Garcia et al. 2022; Correa et al. 2021 |
| **Diplopoda** | Detritivores*<br>Omnivores | Plant litter, fungi, algae, invertebrate carrion | Generally considered dependent on mesic environments. However, some taxa excel in xeric habitats. Resource biomass expected to increase with moisture. |  | Potapov et al. 2022; Hoffman & Payne 1969 | Cloudsley-Thompson 1951; Cloudsley-Thompson 1959; Crawford et al. 1987; Bhakat 2014; David 2015 |
| **Embiopoda** | Phycophages<br>Detritivores | Lichen, algae, plant litter | Inhabit diverse environments from rainforest to dry woodland. *Metoligotoma* are the predominant taxa in the study region, and are expected to prefer dry forest. Resources should be constant among habitats; lichen is abundant in dry forest while litter biomass increases towards rainforest. |  | Edgerly et al. 2005; Miranda-González & McCune 2020; Miller & Edgerly 2008; Potapov et al. 2022 | Edgerly et al. 2005; Miller et al. 2012; Edgerly et al. 2020 |
| **Gastropoda** | Microbivores<br>Herbivores<br>Detritivores<br>Omnivores | Living plants, plant litter, microbes, invertebrate carrion, faeces, fungi, lichens, algae, soil | Inhabit diverse environments ranging from xeric to hygric conditions. Capacity to aestivate and prevent desiccation - preferences for moisture vary drastically across taxa. Resource biomass expected to increase with moisture. |  | Wieser 1978; Spieser & Barker 2001; Potapov et al. 2022 | Schweizer et al. 2019; Hettnerbergerová et al. 2013; Moreno-Rueda et al. 2009; Książkiewicz-Parulska & Ablett 2017 |
| **Isopoda** | Detritivores* | Plant litter, rotting wood, invertebrate carrion | Generally dependent on hygric conditions. However, habitat preferences are diverse with some lineages adapted to xeric conditions. Resource biomass expected to increase with moisture. |  | Wieser 1978; Potapov et al. 2022 | Vilicsics et al. 2005; Lindqvist 1968; Warburg 1964; Leclercq-Dansart et al. 2019; Warburg et al. 1978; Warburg & Berkovitz 1978 |
| **Onychophora** | Carnivores* | Invertebrates | Expected to prefer high moisture environments and have high rates of water loss relative to body size. Prey biomass expected to increase with moisture. |  | Haritos et al. 2010; Read & Hughes 1987; | Reinhard & Rowell 2005; Haritos et al. 2010; Clusella-Trullas & Chown 2008 |
| **Opiliones** | Carnivores*<br>Omnivores<br>Detritivores | Invertebrates, fungi, fruit, decaying fruit, invertebrate carrion | Expected to prefer high moisture environments. Resource biomass expected to increase with moisture. |  | Agosta & Machado 2007; Powell et al. 2021; Potapov et al. 2022; Nyffeler et al. 2023 | Todd 1949; Almeida-Neto et al. 2006 |
| **Pseudoscorpiones** | Carnivores* | Small invertebrates | Data lacking on moisture preferences. Limited evidence suggests no clear preference for moisture. Prey biomass expected to increase with moisture. |  | Garcia et al. 2016; Moura et al. 2018; Potapov et al. 2022 | Villarreal et al. 2019; Adis et al. 1988; Aguiar et al. 2006; Liebke et al. 2021 |

\* = primary trophic role

The forests of south-eastern Australia provide an ideal natural experiment for understanding how invertebrates respond to moisture-graded ecosystems (Figure 1). Across the continent, topographic and geological variation has generated abrupt transitions in soil moisture retention and bushfire susceptibility, resulting in mesic fire-resistant rainforest refugia embedded within a matrix of more fire-prone wet and dry schlerophyll forest (Bowman 2000; Fensham et al. 2024). Subsequently Australia’s rainforests have been dubbed “islands of green in a land of fire” (Bowman 2000). This heterogeneous mixture of forests forms a naturally replicated environmental gradient of forest moisture types, with corresponding increases in biomass and primary productivity toward wetter habitats (Keith et al. 2009; Thomas et al. 2014; Aponte et al. 2020). Notably, these contemporary moisture gradients arose from a long evolutionary history of climatic drying. Much of Australia was formerly dominated by mesic vegetation followed by aridification over the last 30 million years which expanded sclerophyll forests and favoured the spread and diversification of many dry-adapted and fire-tolerant lineages – a critical turning point in the diversification of the Australian biota (Bowman et al. 2000; Byrne et al. 2008; Byrne et al. 2011). This provides good reason to suspect that transitions in invertebrate abundances between dry and wet habitat types are likely to reflect not only shifts in productivity and microclimate, but also differing responses among invertebrate lineages with fundamentally different evolutionary adaptations to water balance.

**Figure 1.**
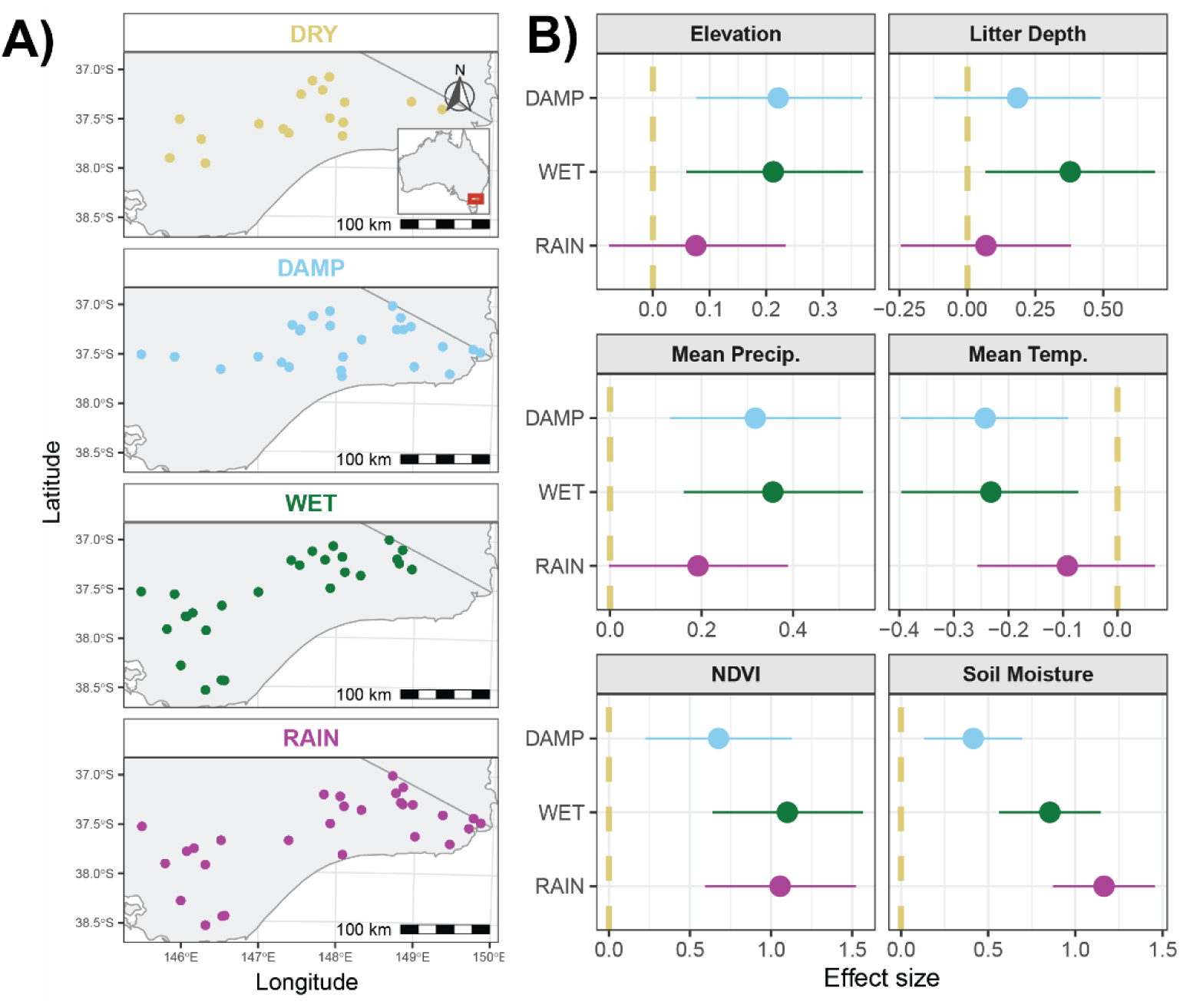
A) Maps of sampling sites. B) Posterior estimates of bioclimatic and environmental variables by habitat type, from separate Bayesian Gaussian models fitted in INLA with habitat type as the sole predictor (dry forest as the reference level; dashed line). Points show posterior means and error bars show 95% credible intervals. Elevation was measured in meters; litter depth was measured in millimetres at three points per quadrat and averaged; mean annual precipitation and mean annual temperature were obtained from the WorldClim database; mean summer NDVI is averaged across 5 years from 16-day, 250 m MODIS composites; soil moisture (% volumetric water content) was measured on site in each quadrat.

Here, we test two alternate hypotheses on leaf litter invertebrate responses to forest moisture gradients: (1) A moisture-productivity hypothesis: where total invertebrate abundance increases from dry sclerophyll forests to rainforest across all taxa; (2) A trait filtering hypothesis: where the abundance of moisture-sensitive orders increase with forest wetness and xeric-adapted taxa show neutral or negative responses, resulting in limited net abundance change underpinned by differences in taxon-specific responses. To do so we quantify leaf litter invertebrate assemblages across 100 sites spanning a wide moisture gradient throughout the south-east forests of Australia. We focus on 11 invertebrate taxa representing broad variation in desiccation tolerance and trophic ecology (Table 1): Isopoda, Gastropoda, Amphipoda, Coleoptera, Diplopoda, Chilopoda, Embioptera, Onychophora, Opiliones, Pseudoscorpiones, and Blattodea. Understanding patterns in general versus taxon-specific abundance responses to moisture will allow us to disentangle the relative contributions of moisture-based productivity and environmental filtering in shaping litter invertebrate communities, with implications for predicting how forest decomposition and nutrient cycling may respond to increasing drought frequency and intensity under climate change.

## Methods

### Vegetation characterisation and site selection

Sites were selected across Victoria (n=100), south eastern Australia (Figure 1A) to represent sclerophyll forests along a moisture gradient (dry, damp, wet) and rainforests at elevations between 5–1350 m. Habitat types were defined using Ecological Vegetation Classes (EVCs) (DEECA 2026, Native Vegetation; Modelled 2005 Ecological Vegetation Classes), which classify vegetation at 1:100,000 resolution based on canopy composition, moisture availability, and plant assemblages, supplemented by a rainforest-specific layer derived from Sentinel imagery at ∼10 m resolution (DEECA 2026, Rainforest Mapping for state-wide Victoria). To target relatively undisturbed habitats, sites were excluded if they had been logged within 20 years (based on the Harvested Logging Coupes dataset; DEECA 2026) or had experienced moderate to severe fire within 10 years (based on Fire Extent and Severity Mapping and Fire History datasets; DEECA 2024, 2026). The Suitability Modeler tool in ArcGIS Pro was used to generate spatial layers representing habitat quality (vegetation type and disturbance history), accessibility (slope <35° derived from Vicmap Elevation 10m DEM and proximity to roads <3 km), with all layers standardized to 10 m resolution and weighted by importance to habitat quality. Sites were manually selected from these suitability maps to represent a range of elevations and habitat types, grouped into replicate blocks of 2–4 habitat types within 10 km of each other to capture ecological variation while minimizing spatial effects. Final locations were confirmed by ground-truthing EVCs to ensure that designated habitat types (dry, damp, wet or rainforest) were correct. In summary, 100 sites were sampled interspersed across the four habitat types, with one transect of ten 0.5 m^2^ leaf litter samples per site.

### Sampling methods

Litter collection was undertaken between September 24^th^, 2024, to December 4^th^, 2024. Sampling blocks (containing at least three categories of the four habitat types: wet, dry, rain, damp) were all sampled on the same date. At each site, we laid out a 90 m transect, always beginning at least 30 m from the nearest road. Ten sampling points were located at 10 m intervals along the transect. Because rainforest primarily occurred as linear strips along narrow, meandering gullies, transects were occasionally bent to remain within rainforest. At each of the 10 sampling points, a quadrat of 50 × 50 cm was placed on the forest floor and several measurements were taken within the quadrat.

### Litter sampling

All leaf litter was collected down to the mineral earth from within each of the 50 × 50 cm quadrats located along the transect (*n* = 10 samples at each site, across 100 sites, for a total of 1000 litter samples). Litter samples were sieved in situ with a 12 mm^2^ sieve to remove coarse material. The leaf litter passing through the sieve was kept in a single pillowcase per quadrat sample, misted with water, and kept cool in plastic storage tubs during transport to a cool room (4 ± 1 °C) (within 5 days after collection). Course debris was discarded in the field. Invertebrates were extracted from the leaf litter within 7 ± 1 days from collection. Leaf litter was processed in 60 cm radius Tullgren funnels with a 70-watt incandescent globe as a light and heat source, and 100% ethanol for specimen capture. Leaf litter was spread across Tullgren funnels to a depth of ≤ 5 cm and heated for 24h. Where all funnels were full, remaining samples were kept in a cool room (4 ± 1 °C) until funnels became available (up to a maximum of 72 h in cold room). Samples (10 quadrats from 100 sites, n = 1000) were sorted by hand using a MZ Leica microscope with 11 taxa counted and transferred to individual tubes in 100% ethanol.

### Bioclimatic variables

The quadrat-level environmental variables were measured on site: 1) Soil moisture (% volumetric water content) was recorded at the centre of each quadrat (n = 10 per site) with a soil moisture meter probe (Fieldscout TDR 150 soil moisture meter probe, Spectrum Technologies). 2) Litter depth was measured with a ruler by taking the average depth across three points within each quadrat (n = 10 per site) (depth was measured from the hard soil layer to the top of litter, in millimetres).

The site-level environmental variables were extracted from remote products: 1) Mean annual precipitation and mean annual temperature were derived from the WorldClim database (Fick & Hijmans 2017) at the geographic coordinates of each site. 2) Elevation for each site was derived from the Vicmap Elevation 10 m Digital Elevation Model (DEECA, 2025), a statewide raster elevation dataset for Victoria, Australia. 3) Vegetation structure/productivity was quantified using the Normalised Difference Vegetation Index (NDVI) accessed from the MODIS remote sensor using the MODIStools package in R, which enables retrieval of MODIS data. We extracted 16-day composite NDVI values at 250 m spatial resolution for each site over a five-year period (2018-2023). To represent site-level vegetation potential, we calculated the mean yearly maximum NDVI value observed across the time series for each site. This metric correlated with overall maximum across a 5-year period and reflects peak canopy greenness and serves as a proxy for maximum primary productivity at each site.

### Statistical Analysis

All statistical analyses were conducted in R (version 4.5.2). Prior to modelling, pairwise Pearson correlations among candidate environmental covariates - mean annual temperature, annual precipitation, elevation, soil moisture, litter depth, and NDVI - were examined to identify collinearity. Correlations were classified as strong (|r| > 0.8), moderate (0.5–0.8), or weak (0.1–0.5). Elevation and mean annual temperature were strongly negatively correlated (r = −0.99) and mean annual precipitation and soil moisture were moderately correlated (r = 0.51). To avoid including redundant predictors, mean annual temperature and annual precipitation were excluded from subsequent models, retaining elevation, soil moisture, litter depth and NDVI as the continuous covariates.

To characterise environmental differences among forest habitat types, Bayesian Gaussian linear models were fitted to each retained environmental variable separately using INLA (Rue et al., 2009) via the inlabru package (Bachl et al., 2019) with habitat type as the sole predictor. Site-level variables (elevation, NDVI, mean annual precipitation, mean annual temperature) were modelled at the site level, while quadrat-level variables (soil moisture, litter depth) included a site-level random intercept to account for repeated measurements within sites. A Gaussian spatial random field (SRF; see below) was included in each environmental model to account for spatial autocorrelation in the environmental variables across the landscape.

To understand how invertebrate abundance changed across habitat types and with environmental covariates, data were modelled using Bayesian negative binomial regression implemented via INLA (Rue et al., 2009) with the inlabru package (Bachl et al., 2019). A negative binomial likelihood was selected to account for overdispersion commonly observed in count data. Analyses were conducted in two stages. First, abundance records were summed across all taxa within each sample to produce a total invertebrate abundance, which was modelled to characterise broad community-level responses to habitat and environmental gradients. Second, the same approach was applied separately to each taxon to examine taxon-specific responses.

Invertebrate abundance was modelled as a function of forest habitat type (dry, damp, wet forest, and rainforest), litter depth, soil moisture, elevation, and NDVI. Habitat type was included as a fixed effect using factor contrast coding, with dry sclerophyll forest as the reference level. The four continuous covariates were each modelled as nonlinear smooth functions using second-order random walks (RW2). Smoothness for each RW2 term was controlled by a penalised complexity prior on the precision parameter (P(σ > 2) = 0.05). A site-level random intercept was included to account for repeated sampling (n = 10 quadrats) within sites, also assigned a PC prior (P(σ > 2) = 0.05).

To account for residual spatial autocorrelation in abundance not explained by the environmental covariates, a Gaussian spatial random field (SRF) was included in each model, implemented as a Matérn covariance structure via the SPDE approach of Lindgren et al. (2011). A triangulated mesh was constructed over the study area using a non-convex hull around sampling locations, with a maximum interior edge length of 8 km. PC priors were placed on the Matérn range (P(range < 80 km) = 0.05) and marginal standard deviation (P(σ > 2) = 0.05) parameters. All models were fitted using a Laplace approximation strategy with the CCD integration scheme.

Posterior distributions of habitat effects were summarised by their posterior mean and 95% credible intervals. To facilitate direct comparison between all habitat types - including the reference level - all pairwise contrasts between habitat types were computed by drawing 1000 posterior samples from the fitted model using inla.posterior.sample() and calculating pairwise differences in the habitat effects on the log abundance scale (Supplementary Figure 1). A contrast whose 95% credible interval excludes zero provides evidence of a difference in abundance between the two habitat types. The nonlinear effects of continuous covariates were visualised separately by predicting the marginal posterior of each RW2 smooth across the observed range of that covariate, with all other model terms held constant.

## Results

### Forest habitats differed in environmental characteristics

Environmental characteristics varied across habitat types (Figure 1B). Damp and wet forests tended to occur at higher elevations, while dry and rainforest sites occurred at lower and overlapping elevations. Mean annual precipitation was substantially lower in dry forests, with all other habitat types experiencing higher and broadly similar rainfall, although rainforest sites showed only marginally higher values relative to dry forest. Mean annual temperature closely tracked elevation, being lower in damp and wet forests and similar between dry and rainforest sites. NDVI was lowest in dry forest, whereas damp, wet, and rainforest forests exhibited comparable values. Litter depth was broadly similar across habitats, except for higher values in wet forest relative to dry forest. Soil moisture showed the clearest separation among habitat types, with dry forests exhibiting the lowest values; damp, wet, and rainforest forests overlapped partially, although damp forest soil moisture was significantly lower than rainforest (Figure 1B), supporting the usefulness of these habitat categories to test the moisture-productivity hypothesis.

### Total invertebrate abundance increased substantially from dry forest to rainforest

A total of 45,050 specimens were sorted and counted (8,491 Amphipoda, 319 Blattodea, 2447 Chilopoda, 20,270 Coleoptera, 4,387 Diplopoda, 66 Embioptera, 1016 Gastropoda, 5772 isopoda, 91 Onychophora, 636 Opiliones, 1555 Pseudoscorpiones).

Total invertebrate abundance was clearly different between habitat types (Figures 2–4) with total invertebrate abundance increasing threefold from dry forest to rainforest, supporting the moisture-productivity hypothesis. Dry forests supported the lowest abundance, significantly lower than all other habitat types. Rainforest sites had the highest abundance, exceeding damp forests and showing a marginal increase relative to wet forests (Figures 2). Wet and damp forests did not differ significantly. Relationships between total invertebrate abundance and environmental covariates revealed consistent but nonlinear responses (Figure 3–6). The isolated effect of litter depth showed a positive monotonic relationship with total invertebrate abundance (Figure 6). The isolated effect of soil moisture exhibited a unimodal response, peaking at approximately 40% volumetric water content and declining sharply above ∼60% (Figure 5). Elevation had no detectable effect on total abundance (Supplementary Figure 2). NDVI showed a threshold response, with lower abundance at maximum NDVI values below ∼0.83, after which abundance plateaued (Supplementary Figure 3).

**Figure 2.**
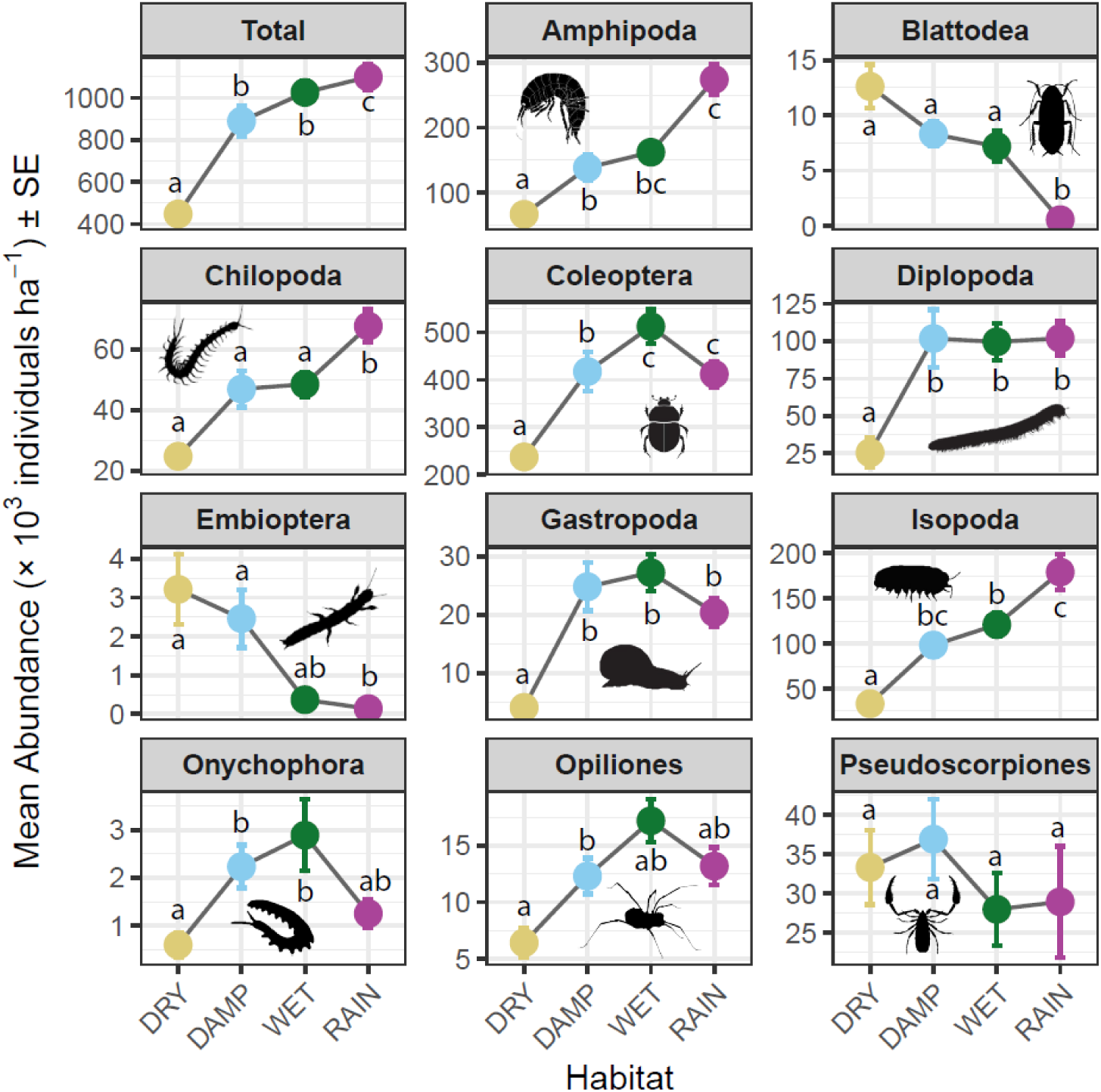
Mean invertebrate abundance (± SE) per hectare of litter across habitat types, estimated by scaling raw counts from 0.5 m² litter samples. Habitats not sharing a letter show non-overlapping 95% credible intervals for their pairwise contrast, based on the Bayesian INLA model (Supplementary Figure 1) and thus differ statistically.

**Figure 3.**
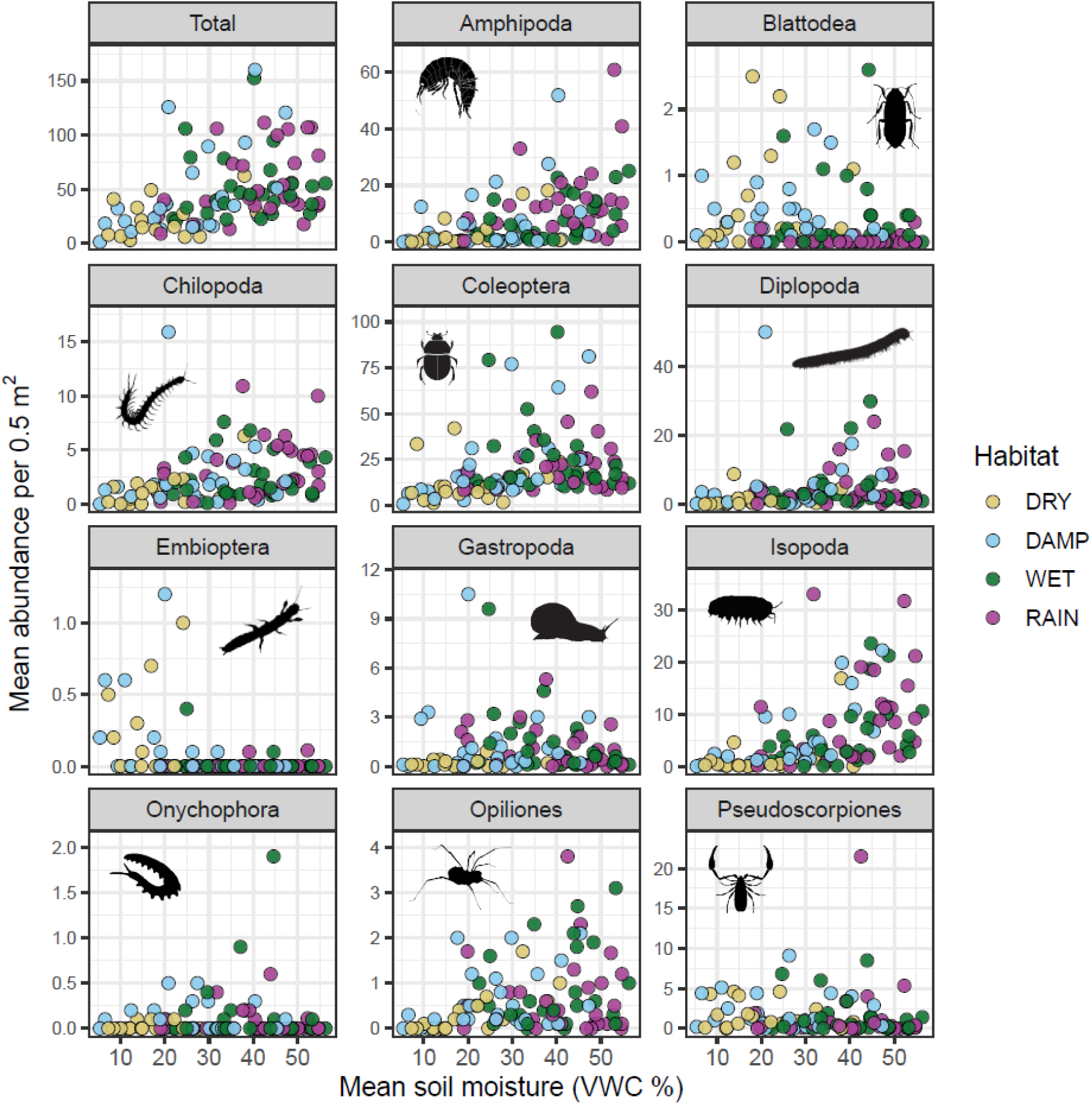
The relationship between mean soil moisture (% volumetric water content) and mean invertebrate abundance at each site. Mean soil moisture was calculated by averaging soil moisture measured in the ten 0.5 m^2^ sampling plots at each site. Mean abundance was calculated by averaging the abundance of all invertebrates, or each taxon, across the same ten 0.5 m^2^ sampling plots at each site. Points are coloured by habitat type.

**Figure 4.**
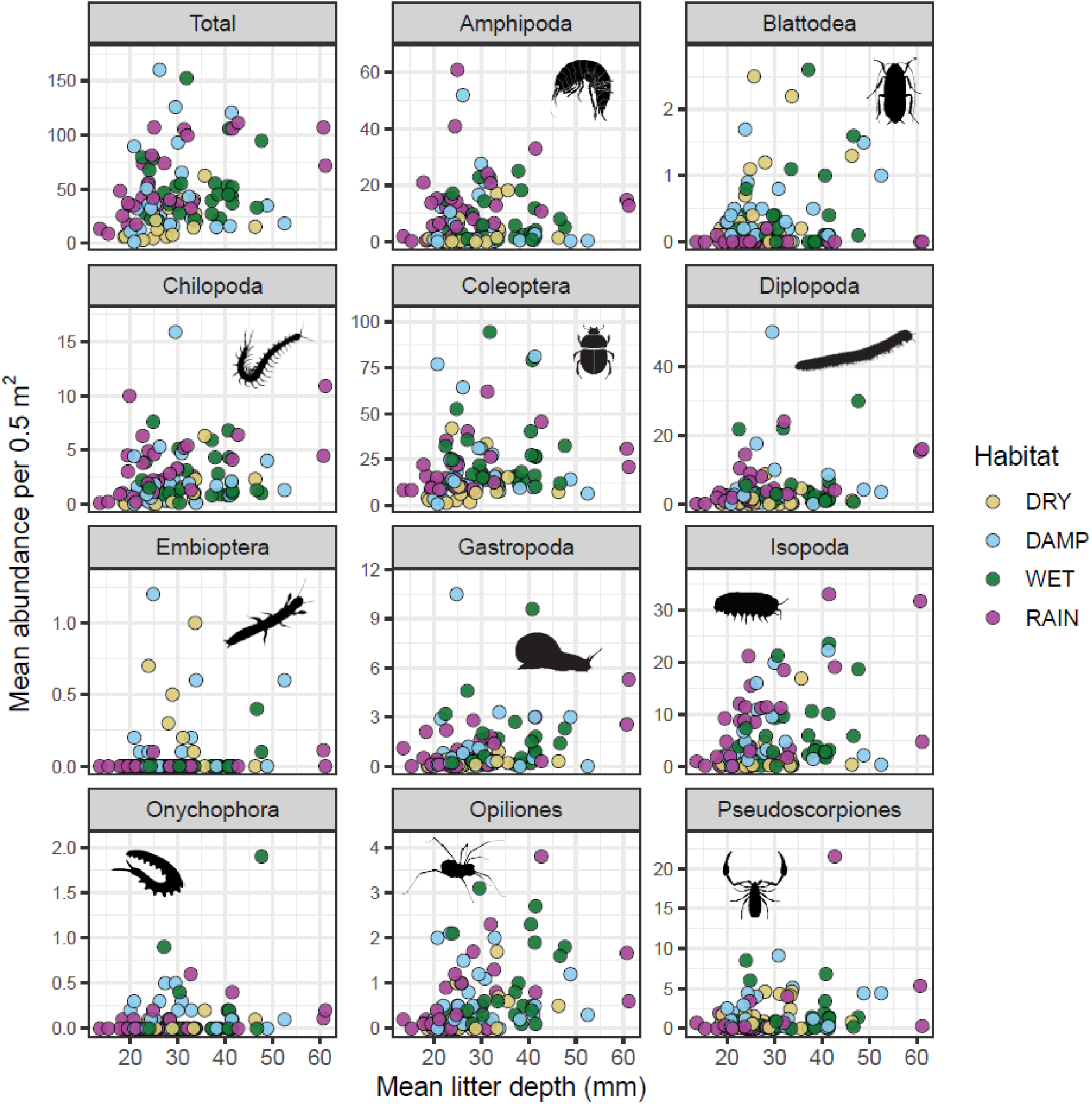
The relationship between mean litter depth (mm) and mean invertebrate abundance at each site. Mean litter depth was calculated by averaging the litter depth measured in the ten 0.5 m^2^ sampling plots at each site. Mean abundance was calculated by averaging the abundance of all invertebrates, or each taxon, across the same ten 0.5 m^2^ sampling plots at each site. Points are coloured by habitat type.

**Figure 5.**
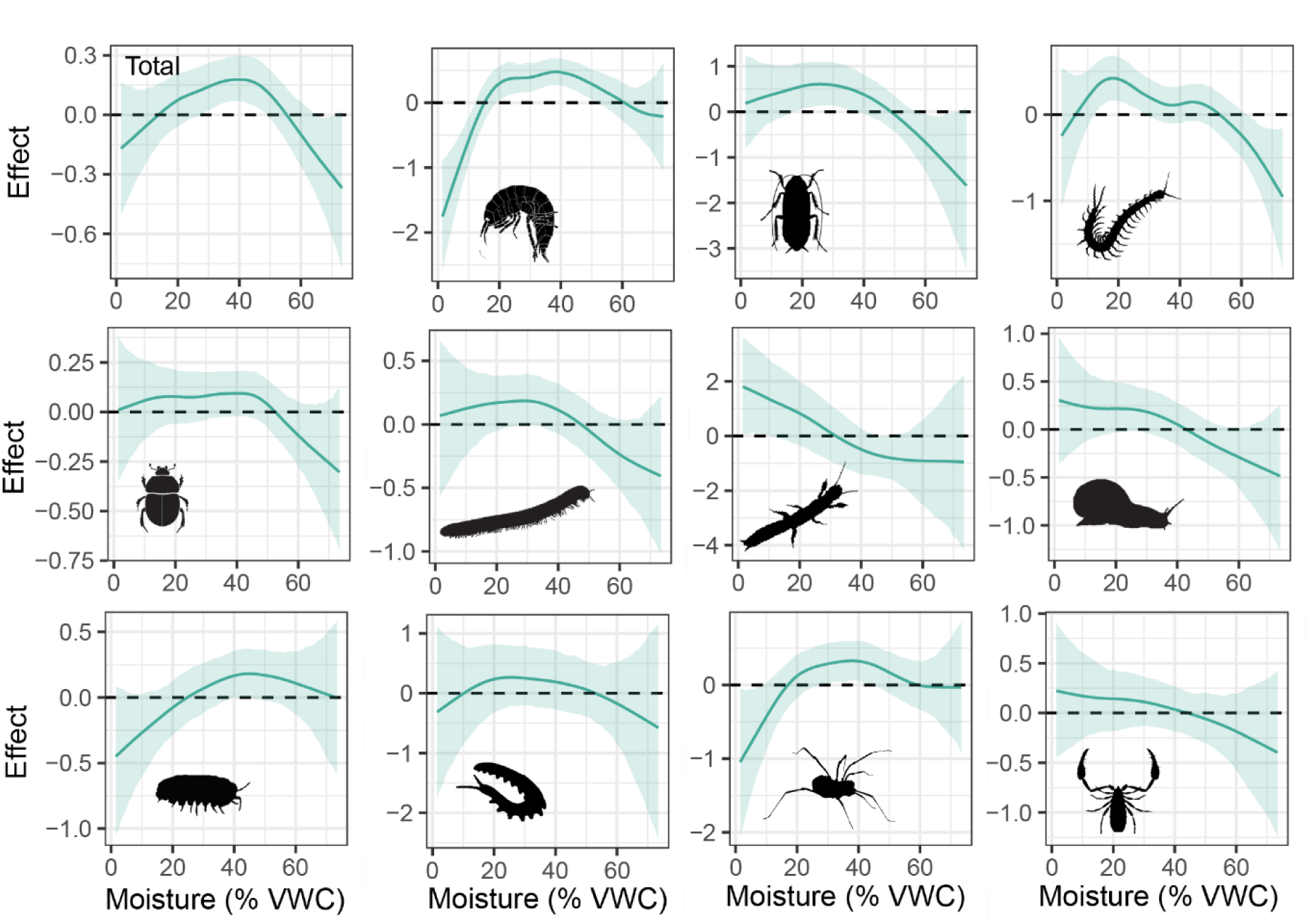
Estimated effect of soil moisture (% volumetric water content) on invertebrate abundance for total invertebrates and each of the 11 taxonomic orders, from the Bayesian INLA model. Solid lines show the model’s estimated relationship between moisture and abundance; shaded ribbons show the 95% credible interval (uncertainty range) around that estimate. Values above zero indicate higher-than-average abundance at that moisture value, and values below zero indicate lower-than-average abundance (log scale).

**Figure 6.**
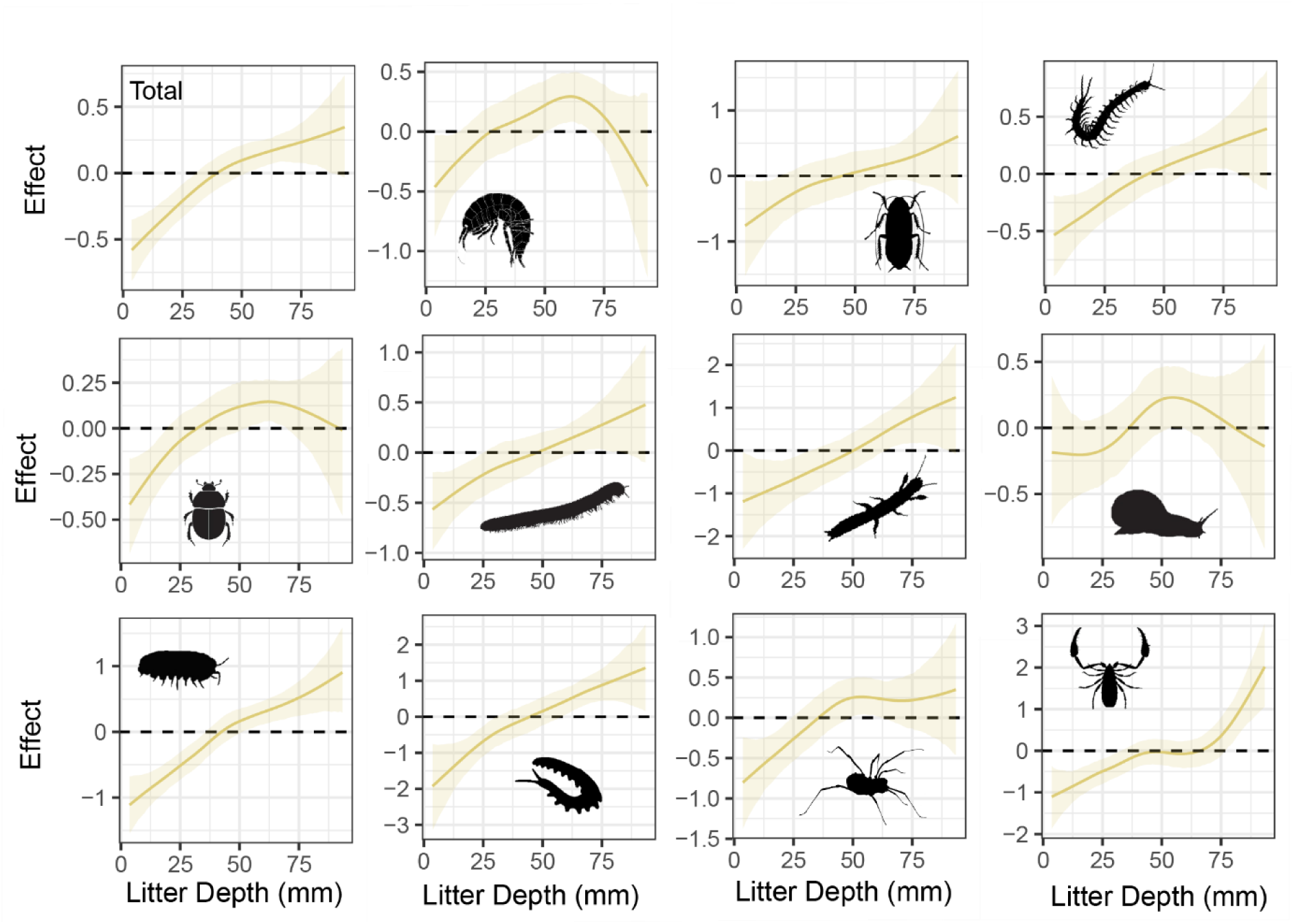
Estimated effect of litter depth (mm) on invertebrate abundance for total invertebrates and each of the 11 taxonomic orders, from the Bayesian INLA model. Solid lines show the model’s estimated relationship between litter depth and abundance; shaded ribbons show the 95% credible interval (uncertainty range) around that estimate. Values above zero indicate higher-than-average abundance at that litter depth value, and values below zero indicate lower-than-average abundance (log scale).

### Invertebrate orders show distinct moisture-associated habitat affinities

Taxon-specific responses to habitat type were pronounced (Figure 2-4), clearly supporting the trait filtering hypothesis. Several taxa (Amphipoda, Coleoptera, Diplopoda, Gastropoda) showed strong reductions in dry forests, with statistically higher abundances in wetter habitats (Figures 2–4; Supplementary Figure 1). In particular, Isopoda, and Chilopoda exhibited strong reductions in dry forest and increases in rainforest exceeding those in wet forests. Other taxa (Onychophora, Opiliones, Pseudoscorpiones) showed weak or inconsistent habitat responses, with relatively uniform abundances across moisture gradients. Blattodea and Embioptera indicated a preference for drier conditions, with higher abundances in dry forests and reduced occurrence in wetter and rainforest habitats contrasting with the moisture responses of other taxa.

### Taxa-specific nonlinear responses to environmental gradients

Responses to environmental gradients varied among taxa (Figures 5–6; Supplementary Figures 2-3). Several taxa (Coleoptera, Diplopoda, Gastropoda, Onychophora, Pseudoscorpiones) showed weak or negligible responses to soil moisture (Figure 5), whereas Amphipoda, Isopoda, Chilopoda, Opiliones, and Blattodea exhibited strong positive responses to increasing moisture. Embioptera declined with increasing moisture. Across most taxa, abundance declined at the highest moisture levels (>40–50% VWC), with exceptions among the most moisture-adapted groups (Amphipoda, Isopoda, Opiliones), which maintained higher abundances under wet conditions. All taxa except Gastropoda showed positive associations with litter depth (Figure 6). Elevation effects were limited, with significant responses detected only for Coleoptera, Amphipoda, and Chilopoda (Supplementary Figure 2). Amphipoda peaked at intermediate elevations (∼500 m) and declined above ∼1100 m, Coleoptera peaked near ∼1000 m, while Chilopoda increased monotonically with elevation. NDVI effects (Supplementary Figure 3) were detected in Amphipoda, Blattodea, Chilopoda, Opiliones, Pseudoscorpiones, and Isopoda, generally showing increased abundance at intermediate to high NDVI, except Pseudoscorpiones, which declined at high NDVI.

## Discussion

Moisture availability structures the patterns of biomass and productivity of the world’s forests (Silvertown et al. 1994; Keith et al. 2009; Thomas et al. 2014; Pan et al. 2013; Malhi et al. 2015; Moore et al. 2018; Kicklighter et al. 1999; Aponte et al. 2020; Muller-Landau et al. 2021). Yet how these moisture gradients shape the detrital communities responsible for maintaining forest nutrient cycling remains poorly understood. Here, we show that litter invertebrate abundance increases threefold across habitat types from dry sclerophyll forest to rainforest, thus linking detrital communities to the same broad moisture–productivity relationships that shape plant communities (Silvertown et al. 1994; Keith et al. 2009; Thomas et al. 2014; Pan et al. 2013; Malhi et al. 2015; Moore et al. 2018; Kicklighter et al. 1999; Aponte et al. 2020 et al. 2015; Muller-Landau et al. 2021). However, we did not observe simple linear responses to soil water availability. Instead, invertebrate abundance peaked at intermediate soil moisture levels, and individual orders showed positive, weak, or negative responses across the moisture gradient. Together, these findings demonstrate the role of two mechanisms that generate distinct assemblages and population densities across habitat types – a broad moisture-productivity effect and an environmental/trait filtering effect.

We found that wetter forests supported greater total invertebrate abundance supporting the moisture-productivity hypothesis. This is not unexpected, as it is well appreciated that relative to dry forests, wetter forests have higher plant biomass, litterfall rates, and litter decomposition rates (Murphy et al. 1998; Cisneros-Dozal et al. 2007; Zhang et al. 2008; Thomas et al. 2014; Muller-Landau et al. 2021; Fensham et al. 2024) – all of which should generally increase the abundance of litter invertebrates. With the exception of Gastropoda, invertebrate abundance across all orders was also positively associated with litter availability, reinforcing the idea that litter invertebrate communities are closely coupled to rates of organic matter cycling across forest moisture gradients. Higher rates of decomposition under wetter conditions may explain why rainforest sites contained relatively shallow litter layers in our data (Figure 1B) despite supporting the highest abundances of litter fauna – organic matter may be processed more rapidly and consistently in rainforests rather than accumulated. Australian rainforest communities have historically been considered to produce leaves of higher nutrient quality (relative to the recalcitrant litter of *Eucalyptus* which dominate dryer habitat types) – though evidence is conflicting (Bowman 2000; Parsons & Congdon 2008; Thomas et al. 2014; Fensham et al. 2024). In contrast, wet sclerophyll forests were the only forests that had deeper litter layers relative to dry forests (Figure 1B) yet slightly lower total invertebrate abundances compared to rainforests, possibly reflecting the slower decomposition rates of lignified recalcitrant sclerophyllous litter, which may have lower energetic and nutrient availability, despite higher density (Specht & Rundle 1990; Talbot et al. 2012; Thomas et al. 2014).

Beyond total invertebrate abundance, strong variation in the composition of major taxonomic groups across the habitat gradient revealed that trait filtering reorganizes detrital communities rather than simply reducing their abundance under drier conditions. Moisture-sensitive groups – notably Isopoda and Chilopoda – were concentrated in rainforest habitats, and Amphipoda, Coleoptera, Diplopoda, and Gastropoda all declined sharply in dry forests. However, Opiliones, Onychophora, and Pseudoscorpiones showed weak relationships with forest moisture gradients, and Blattodea and Embioptera reached peak abundances in dry sclerophyll forests and declined sharply toward rainforests. That these lineage-specific responses to forest moisture type emerged at such high taxonomic levels reinforces the notion that deeply conserved differences in water balance strategies (Hurley 1968; Edney 1968; Schweizer et al. 2019; Warburg 1965) are likely determinants of community assembly across forest moisture gradients. Whether equivalent compositional changes occur at finer taxonomic scales, including among species within these orders, is likely however as even the most moisture-sensitive groups such as Amphipoda had representatives capable of persisting in relatively dry habitats (22 individuals were collected from a single 0.5 m^2^ sample from a dry forest at only 14.4% soil moisture), possibly representing distinct species or populations from those dominating rainforest assemblages. In fact, studies show that dry forests, wet forests, and rainforests can have distinct litter invertebrate assemblages even when separated by small spatial scales (González & Seastedt 2000; Houston & Melzer 2012). Considering that dry forests in eastern Australia are evolutionarily younger than wet forests, it is particularly likely that many taxa have diversified into dry forest habitat over the last 30 million years following Cenozoic aridification of the continent (Byrne et al. 2008; Byrne et al. 2011).

With increasing climate change-driven aridity and fire risk, wet forests are often considered the ecosystems most at risk. However, our data also support the characterisation of dry forests as distinctive and ecologically significant habitats for Australian litter invertebrates. Australia’s dry forests have been comparatively neglected in invertebrate conservation research despite growing recognition of the importance of dry forests to global biodiversity (Kiley et al. 2017; Janzen et al. 1988; Bastin et al. 2017; Murphy & Lugo 1986; Miles et al. 2006; Houston & Melzer 2012) and the likelihood that forest communities adapted to xeric conditions may nonetheless be approaching the limits of their adaptive capacity under increasing drought frequency (Aguirre-Gutiérrez et al. 2019; Miles et al. 2006). Although invertebrate abundances in dry forests were lower than in wetter forests, dry forests still supported substantially productive assemblages – approximately 400,000 individuals per hectare (roughly one-third of rainforest densities), including ∼70,000 amphipods and ∼30,000 isopods per hectare. Critically, these dry-forest communities were not simply impoverished versions of wetter-forest assemblages but were compositionally distinctive, structured by increased abundances of Embioptera and Blattodea suggesting that these taxa are specialised for the more xeric conditions of dry schlerophyll forest. The Embioptera in our dry forest samples belong predominantly to the genus *Metoligotoma* which are endemic to Australia and construct silken galleries within surface litter and thus depend on dried leaf material both as shelter substrate and dietary resource (Davis 1938, 1942, 1943; Miller & Edgerly 2008). Similarly, many lineages of Australian cockroaches (Blattodea) are known to have diversified into xeric habitats following historical contractions of mesic biomes and exhibit physiological and behavioural adaptations to low-moisture environments (Bell et al. 2007; Lo et al. 2016). Whether these compositionally distinctive communities of dry forests also differ functionally in rates of per-capita litter fragmentation or decomposition efficiency remains an important knowledge gap for predicting how nutrient cycling will respond to increasing drought frequency. Evidence from both terrestrial and aquatic litter studies suggests that the functional composition of litter-feeding invertebrate communities can influence decomposition rates (Heemsbergen et al. 2004; McKie et al. 2008; Zimmer et al. 2005; Jonsson & Malmqvist 2003; McKie et al. 2009; Gessner et al. 2010; del Campo et al. 2025) and that invertebrates play disproportionately important roles in nutrient cycling within xeric habitats (Ward & Seely 1996; Cheli et al. 2022; Seely & Luow 1980; Silva et al. 1985; Cepeda-Pizarro & Whitford 1989; Njoroge et al. 2021; Sagi & Hawlena 2023; Torsekar et al. 2024). Taken together, these considerations make the functional characterisation of dry-forest detrital communities an important avenue for future research.

Overall, our findings demonstrate that forest moisture gradients simultaneously regulate the abundance and taxonomic composition of litter communities. Two questions follow most directly from these results. First, under aridification and drought, do invertebrate communities in wet forests transition toward assemblages characteristic of drier systems, or simply persist as numerically diminished wet forest assemblages? We only consider spatial variation in moisture gradients here, but artificial manipulation of moisture in forest plots, or comparison of functionality across El Niño and La Niña periods would reveal temporal trends (as per Chikoski et al. 2006; Landesman et al. 2011). Second, do changes in community composition along these moisture gradients alter the functional performance of the detrital system – for example, do dry forest taxa process dry litter at different rates than their counterparts in wet forests and vice-versa? Addressing these questions would require transplanting litter and invertebrate fauna between the different habitat types, combined with measurements of decomposition function and community composition (following the design of Joly et al. 2023). Determining if similar moisture–community relationships occur across forest biomes globally, whether compositionally distinct detrital communities are functionally interchangeable, and how drying forest shifts in community structure, will be essential knowledge to determine the resilience of decomposition and nutrient cycling pathways as forest ecosystems face increasing drought worldwide.

## Acknowledgements

We thank Andrew Chalmers, Albin Larsson Ekström, Finlay Davidson, Andrejs Enriquez, Dennis Black, Erica Dudek, Colin Moffa, Tara Pridham, Eva Billman, Lucas Dunipace, Anita Thamm, Nicholas Gale, Nicolas Wollek, Kate Kearney, Paige Matheson, Charlie Disher, Aidan Fitt, Daniel Kurek, Alex Bester-Lowe, Angie Symon, Maya Manganaro, Anya Smolders, Eboney Hazel Teuber, Elijah Kune, Emerson Taylor Laherty, Randy Marks, and Amelia Corboy for their assistance with fieldwork and sorting specimens. Field collections were undertaken under the approval of the Victorian Government Department of Energy, Environment and Climate Action (DEECA) in line with the Wildlife Act 1975. Research authorisation permit number 10011226. Research funding for this project was from the Australian Research Council, grant number LP230100176.

## Data availability statement

Data and code are publicly available from the figshare repository https://doi.org/10.6084/m9.figshare.33919279

## Notes

https://doi.org/10.6084/m9.figshare.33919279

### Competing Interest Statement

The authors have declared no competing interest.

## References

Acosta LE, Machado G. 8 diet and foraging. In: Harvestmen: The Biology of Opiliones. Ricardo Pinto-da-Rocha, Glauco Machado, Gonzalo Giribet (eds). Harvard University Press. p 309–338.

Adis J, Mahnert V, de Morais JW, José MGR. 1988. Adaptation of an amazonian pseudoscorpion (arachnida) from dryland forests to inundation forests. Ecology. 69:287–291

Aguiar NO, Gualberto TL, Franklin E. 2006. A medium-spatial scale distribution pattern of pseudoscorpionida (arachnida) in a gradient of topography (altitude and inclination), soil factors, and litter in a central amazonia forest reserve, brazil. Brazilian Journal of Biology. 66:791–802

Aguirre-Gutiérrez J, Oliveras I, Rifai S, Fauset S, Adu-Bredu S, Affum-Baffoe K, Baker TR, Feldpausch TR, Gvozdevaite A, Hubau W. 2019. Drier tropical forests are susceptible to functional changes in response to a long-term drought. Ecology Letters. 22:855–865

Ahearn GA. 1970. The control of water loss in desert tenebrionid beetles. Journal of Experimental Biology. 53:573–595

Allen CD, Breshears DD, McDowell NG. 2015. On underestimation of global vulnerability to tree mortality and forest die-off from hotter drought in the anthropocene. Ecosphere. 6:art129

Allison SD, Lu Y, Weihe C, Goulden ML, Martiny AC, Treseder KK, Martiny JBH. 2013. Microbial abundance and composition influence litter decomposition response to environmental change. Ecology. 94:714–725

Almeida-Neto M, Machado G, Pinto-da-Rocha R, Giaretta AA. 2006. Harvestman (arachnida: Opiliones) species distribution along three neotropical elevational gradients: An alternative rescue effect to explain rapoport’s rule? Journal of Biogeography. 33:361–375

Anderson JT, Smith LM. 2000. Invertebrate response to moist-soil management of playa wetlands. Ecological Applications. 10:550–558

Andriuzzi WS, Franco ALC, Ankrom KE, Cui S, de Tomasel CM, Guan P, Gherardi LA, Sala OE, Wall DH. 2020. Body size structure of soil fauna along geographic and temporal gradients of precipitation in grasslands. Soil Biology and Biochemistry. 140:107638

Aponte C, Kasel S, Nitschke CR, Tanase MA, Vickers H, Parker L, Fedrigo M, Kohout M, Ruiz-Benito P, Zavala MA, Bennett LT. 2020. Structural diversity underpins carbon storage in australian temperate forests. Global Ecology and Biogeography. 29:789–802

Appel AG, Reierson DA, Rust MK. 1983. Comparative water relations and temperature sensitivity of cockroaches. Comparative Biochemistry and Physiology Part A: Physiology. 74:357–361

Bachl FE, Lindgren F, Borchers DL, Illian JB. 2019. Inlabru: An r package for bayesian spatial modelling from ecological survey data. Methods in Ecology and Evolution. 10:760–766

Bastin J-F, Berrahmouni N, Grainger A, Maniatis D, Mollicone D, Moore R, Patriarca C, Picard N, Sparrow B, Abraham EM, Aloui K, Atesoglu A, Attore F, Bassüllü Ç, Bey A, Garzuglia M, García-Montero LG, Groot N, Guerin G, Laestadius L, Lowe AJ, Mamane B, Marchi G, Patterson P, Rezende M, Ricci S, Salcedo I, Diaz AS-P, Stolle F, Surappaeva V, Castro R. 2017. The extent of forest in dryland biomes. Science. 356:635–638

Bell WJ, Roth LM, Nalepa CA. 2007. Cockroaches: Ecology, behavior, and natural history. Johns Hopkins University Press.

Bennett AC, Penman TD, Arndt SK, Roxburgh SH, Bennett LT. 2020. Climate more important than soils for predicting forest biomass at the continental scale. Ecography. 43:1692–1705

Bhakat S. 2014. Comparative water relations of some tropical millipedes. Kragujevac Journal of Science. 36:185–194

Blower JG. 1955. Millipedes and centipedes as soil animals. In Soil zoology: 138–151. D. K. McE. Kevan (Ed.) London: Butterworths.

Bowman DM. 2000a. Australian rainforests: Islands of green in a land of fire. Cambridge University Press.

Bowman DMJS. 2000b. Rainforests and flame forests: The great australian forest dichotomy. Australian Geographical Studies. 38:327–331

Brockett BFT, Prescott CE, Grayston SJ. 2012. Soil moisture is the major factor influencing microbial community structure and enzyme activities across seven biogeoclimatic zones in western canada. Soil Biology and Biochemistry. 44:9–20

Butler OM, Lewis T, Rezaei Rashti M, Maunsell SC, Elser JJ, Chen C. 2019. The stoichiometric legacy of fire regime regulates the roles of micro-organisms and invertebrates in decomposition. Ecology. 100:e02732

Byrne M, Steane DA, Joseph L, Yeates DK, Jordan GJ, Crayn D, Aplin K, Cantrill DJ, Cook LG, Crisp MD, Keogh JS, Melville J, Moritz C, Porch N, Sniderman JMK, Sunnucks P, Weston PH. 2011. Decline of a biome: Evolution, contraction, fragmentation, extinction and invasion of the australian mesic zone biota. Journal of Biogeography. 38:1635–1656

Byrne M, Yeates DK, Joseph L, Kearney M, Bowler J, Williams MAJ, Cooper S, Donnellan SC, Keogh JS, Leys R, Melville J, Murphy DJ, Porch N, Wyrwoll KH. 2008. Birth of a biome: Insights into the assembly and maintenance of the australian arid zone biota. Molecular Ecology. 17:4398–4417

Callaghan CT, Santini L, Spake R, Bowler DE. 2024. Population abundance estimates in conservation and biodiversity research. Trends in Ecology & Evolution. 39:515–523

Carbone C, Rowcliffe JM, Cowlishaw G, Isaac NJB 2007. The scaling of abundance in consumers and their resources: Implications for the energy equivalence rule. The American Naturalist. 170:479–484

Cepeda-Pizarro JG, Whitford WG. 1990. Decomposition patterns of surface leaf litter of six plant species along a chihuahuan desert watershed. The American Midland Naturalist. 123:319–330

Cheli GH, Tomas B, Gustavo EF. 2022. The role of *Nyctelia dorsata* Fairmaire, 1905 (Coleoptera: Tenebrionidae) on litter fragmentation processes and soil biogeochemical cycles in arid patagonia. Annales Zoologici. 72:129–134

Chikoski JM, Ferguson SH, Meyer L. 2006. Effects of water addition on soil arthropods and soil characteristics in a precipitation-limited environment. Acta Oecologica. 30:203–211

Chown SL, Nicolson SW. 2004. Insect physiological ecology: Mechanisms and patterns. Oxford University Press.

Cisneros-Dozal LM, Trumbore SE, Hanson PJ. 2007. Effect of moisture on leaf litter decomposition and its contribution to soil respiration in a temperate forest. Journal of Geophysical Research: Biogeosciences. 112

Cloudsley-Thompson JL. 1959. Microclimate, diurnal rhythms and the conquest of the land by arthropods. International Journal of Bioclimatology and Biometeorology. 3:105–118

Cloudsley-Thompson JL. 1951. On the responses to environmental stimuli, and the sensory physiology of millipedes (Diplopoda). Proceedings of the Zoological Society of London. 121:253–277

Clusella-Trullas S, Chown SL. 2008. Investigating onychophoran gas exchange and water balance as a means to inform current controversies in arthropod physiology. Journal of Experimental Biology. 211:3139–3146

Coleine C, Delgado-Baquerizo M, DiRuggiero J, Guirado E, Harfouche AL, Perez-Fernandez C, Singh BK, Selbmann L, Egidi E. 2024. Dryland microbiomes reveal community adaptations to desertification and climate change. The ISME Journal. 18:wrae056

Collison EJ, Riutta T, Slade EM. 2013. Macrofauna assemblage composition and soil moisture interact to affect soil ecosystem functions. Acta Oecologica. 47:30–36

Correa CMA, da Silva PG, Puker A, Gil RL, Ferreira KR. 2021. Rainfall seasonality drives the spatiotemporal patterns of dung beetles in amazonian forests in the arc of deforestation. Journal of Insect Conservation. 25:453–463

Crawford CS, Bercovitz K, Warburg MR. 1987. Regional environments, life-history patterns, and habitat use of spirostreptid millipedes in arid regions. Zoological Journal of the Linnean Society. 89:63–88

Crowson RA. 1986. The biology of the Coleoptera. Academic press.

David J-F. 2015. 12 Diplopoda—ecology. In: Treatise on zoology-anatomy, taxonomy, biology the Myriapoda, volume 2. Minelli, A (ed). Brill. p 303–327.

David JF. 2014. The role of litter-feeding macroarthropods in decomposition processes: A reappraisal of common views. Soil Biology and Biochemistry. 76:109–118

Davis C. Studies in australian Embioptera. Part iii: Revision of the genus Metoligotoma, with descriptions of new species, and other notes on the family Oligotomidae. In: Proceedings of the Linnean Society of New South Wales. Vol. 63 p 226–272.

Davis C. 1942. Studies in australian Embioptera. Part v. Geographical variation in Metoligotoma reducta. Proc Linn Soc New South Wales. 67:331–334

Davis C. 1943. Studies in australian Embioptera. Part vi. Records of the genus Metoligotoma from victoria. Proc Linn Soc New South Wales. 68:65–66

Davis CA, Austin JE, Buhl DA. 2006. Factors influencing soil invertebrate communities in riparian grasslands of the central platte river floodplain. Wetlands. 26:438–454

de Groot MD, Santamaria B, Adriaens T, Cottrell TE, Maes D, Sakaki S, Verbeken A, Nedvěd O, Haelewaters D. 2026. Effects of temperature and humidity on the presence and prevalence of a common fungal parasite on an invasive ladybird. Ecological Entomology. 51:86–97

del Campo R, Blackman RC, Martini J, Fuß T, Thuile Bistarelli L, Gessner MO, Altermatt F, Singer G. 2025. Functional macroinvertebrate diversity stabilizes decomposition among leaf litter resources across a river network. Ecological Monographs. 95:e70010

Denissen JMC, Teuling AJ, Pitman AJ, Koirala S, Migliavacca M, Li W, Reichstein M, Winkler AJ, Zhan C, Orth R. 2022. Widespread shift from ecosystem energy to water limitation with climate change. Nature Climate Change. 12:677–684

Dunlop JA, Scholtz G, Selden PA. Water-to-land transitions. In: Minelli A, Boxshall G, Fusco G, editors. Arthropod biology and evolution: Molecules, development, morphology. Springer Berlin Heidelberg. p 417–439.

Edgerly JS, Sandel B, Regoli I, Okolo O. 2020. Silk spinning behavior varies from species-specific to individualistic in Embioptera: Do environmental correlates account for this diversity? Insect Systematics and Diversity. 4:2

Edgerly JS, Tadimalla A, Dahlhoff EP. 2005. Adaptation to thermal stress in lichen-eating webspinners (Embioptera): Habitat choice, domicile construction and the potential role of heat shock proteins. Functional Ecology. 19:255–262

Edney EB. 1968. Transition from water to land in isopod crustaceans. American Zoologist. 8:309–326

Eisenhauer N, Ochoa-Hueso R, Huang Y, Barry KE, Gebler A, Guerra CA, Hines J, Jochum M, Andraczek K, Bucher SF, Buscot F, Ciobanu M, Chen H, Junker R, Lange M, Lehmann A, Rillig M, Römermann C, Ulrich J, Weigelt A, Schmidt A, Türke M. 2023. Ecosystem consequences of invertebrate decline. Current Biology. 33:4538–4547.e4535

Erin CP, Christina JP, Anthony JH, Glauco M, Gregory IH. 2021. Diet, predators, and defensive behaviors of new zealand harvestmen (Opiliones: Neopilionidae). The Journal of Arachnology. 49:122–140

Fensham RJ, Laffineur B, Browning O. 2024. Fuel dynamics and rarity of fire weather reinforce coexistence of rainforest and wet sclerophyll forest. Forest Ecology and Management. 553:121598

Fick SE, Hijmans RJ. 2017. Worldclim 2: New 1-km spatial resolution climate surfaces for global land areas. International Journal of Climatology. 37:4302–4315

Friend JA, Richardson AMM. 1986. Biology of terrestrial amphipods. Annual Review of Entomology. 31:25–48

Friend JA, Richardson AMM. 1977. A preliminary study of niche partition in two tasmanian terrestrial amphipod species. Ecological Bulletins. 24–35

Frith D, Frith C. 1990. Seasonality of litter invertebrate populations in an australian upland tropical rain forest. Biotropica. 22:181–190

Fründ H-C. 1987. Räumliche Verteilung und Koexistenz der Chilopoden in einem Buchen-Altbestand. Pedobiologia. 30:19–30

García-Palacios P, Maestre FT, Kattge J, Wall DH. 2013. Climate and litter quality differently modulate the effects of soil fauna on litter decomposition across biomes. Ecology Letters. 16:1045–1053

Garcia LF, Gonzalez-Gomez JC, Valenzuela-Rojas JC, Tizo-Pedroso E, Lacava M. 2016. Diet composition and prey selectivity of colombian populations of a social pseudoscorpion. Insectes Sociaux. 63:635–640

Gessner MO, Swan CM, Dang CK, McKie BG, Bardgett RD, Wall DH, Hättenschwiler S. 2010. Diversity meets decomposition. Trends in Ecology & Evolution. 25:372–380

González G, Seastedt TR. 2000. Comparison of the abundance and composition of litter fauna in tropical and subalpine forests. Pedobiologia. 44:545–555

Grant JA, Villani MG. 2003. Soil moisture effects on entomopathogenic nematodes. Environmental Entomology. 32:80–87

Gunn DL, Cosway CA. 1938. The temperature and humidity relations of the cockroach: V. Humidity preference. Journal of Experimental Biology. 15:555–563

Günther B, Rall BC, Ferlian O, Scheu S, Eitzinger B. 2014. Variations in prey consumption of centipede predators in forest soils as indicated by molecular gut content analysis. Oikos. 123:1192–1198

Haritos VS, Niranjane A, Weisman S, Trueman HE, Sriskantha A, Sutherland TD. 2010. Harnessing disorder: Onychophorans use highly unstructured proteins, not silks, for prey capture. Proceedings of the Royal Society B: Biological Sciences. 277:3255–3263

Heemsbergen DA, Berg MP, Loreau M, van Hal JR, Faber JH, Verhoef HA. 2004. Biodiversity effects on soil processes explained by interspecific functional dissimilarity. Science. 306:1019–1020

Hettenbergerová E, Horsák M, Chandran R, Hájek M, Zelený D, Dvořáková J. 2013. Patterns of land snail assemblages along a fine-scale moisture gradient. Malacologia. 56:31–42

Hobbs NT. 2024. A general, resource-based explanation for density dependence in populations of large herbivores. Ecological Monographs. 94:e1600

Hoffman RL, Payne JA. 1969. Diplopods as carnivores. Ecology. 50:1096–1098

Homet P, Gómez-Aparicio L, Matías L, Godoy O. 2021. Soil fauna modulates the effect of experimental drought on litter decomposition in forests invaded by an exotic pathogen. Journal of Ecology. 109:2963–2980

Houston WA, Melzer A. 2012. Dry rainforests have a distinct and more diverse assemblage of epigaeic invertebrates than eucalypt woodlands: Implications for ecosystem health monitoring. Pacific Conservation Biology. 18:133–145

Hurley DE. 1968. Transition from water to land in amphipod crustaceans. American Zoologist. 8:327– 353

Janzen DH. 1988. Management of habitat fragments in a tropical dry forest: Growth. Annals of the Missouri Botanical Garden. 75:105–116

Janzen DH, Schoener TW. 1968. Differences in insect abundance and diversity between wetter and drier sites during a tropical dry season. Ecology 49:96–110.

Joly F-X, Coq S, Coulis M, Nahmani J, Hättenschwiler S. 2018. Litter conversion into detritivore faeces reshuffles the quality control over C and N dynamics during decomposition. Functional Ecology. 32:2605–2614

Joly F-X, Scherer-Lorenzen M, Hättenschwiler S. 2023. Resolving the intricate role of climate in litter decomposition. Nature Ecology & Evolution. 7:214–223

Jonsson M, Malmqvist B. 2000. Ecosystem process rate increases with animal species richness: Evidence from leaf-eating, aquatic insects. Oikos. 89:519–523

Kaspari M, Weiser MD. 2000. Ant activity along moisture gradients in a neotropical forest. Biotropica. 32:703–711

Keith H, Mackey BG, Lindenmayer DB. 2009. Re-evaluation of forest biomass carbon stocks and lessons from the world’s most carbon-dense forests. Proceedings of the National Academy of Sciences. 106:11635–11640

Kellermann V, Loeschcke V, Hoffmann AA, Kristensen TN, Fløjgaard C, David JR, Svenning J-C, Overgaard J. 2012. Phylogenetic constraints in key functional traits behind species’ climate niches: Patterns of desiccation and cold resistance across 95 *Drosophila* species. Evolution. 66:3377–3389

Kicklighter DW, Bondeau A, Schloss AL, Kaduk J, McGuire AD, Intercomparison T.P.O.F.T.P.N.M. 1999. Comparing global models of terrestrial net primary productivity (npp): Global pattern and differentiation by major biomes. Global Change Biology. 5:16–24

Kiley HM, Ainsworth GB, van Dongen WFD, Weston MA. 2017. Variation in public perceptions and attitudes towards terrestrial ecosystems. Science of The Total Environment. 590-591:440–451

Klemmer AJ, Wissinger SA, Greig HS, Ostrofsky ML. 2012. Nonlinear effects of consumer density on multiple ecosystem processes. Journal of Animal Ecology. 81:770–780

Kraft NJB, Adler PB, Godoy O, James EC, Fuller S, Levine JM. 2015. Community assembly, coexistence and the environmental filtering metaphor. Functional Ecology. 29:592–599

Książkiewicz-Parulska Z, Ablett JD. 2017. Microspatial distribution of molluscs and response of species to litter moisture, water levels and eutrophication in moist, alkaline ecosystems. Belgian Journal of Zoology. 147

Landesman WJ, Treonis AM, Dighton J. 2011. Effects of a one-year rainfall manipulation on soil nematode abundances and community composition. Pedobiologia. 54:87–91

Lazo-Wasem EA. 1984. Physiological and behavioral ecology of the terrestrial amphipod *Arcitalitrus sylvaticus* (Haswell, 1880). Journal of Crustacean Biology. 4:343–355

Leclercq-Dransart J, Pernin C, Demuynck S, Grumiaux F, Lemière S, Leprêtre A. 2019. Isopod physiological and behavioral responses to drier conditions: An experiment with four species in the context of global warming. European Journal of Soil Biology. 90:22–30

Lensing JR, Todd S, Wise DH. 2005. The impact of altered precipitation on spatial stratification and activity-densities of springtails (collembola) and spiders (araneae). Ecological Entomology. 30:194–200

Levings SC, Windsor DM. 1985. Litter arthropod populations in a tropical deciduous forest: Relationships between years and arthropod groups. Journal of Animal Ecology. 54:61–69

Liebke DF, Harms D, Widyastuti R, Scheu S, Potapov AM. 2021. Impact of rainforest conversion into monoculture plantation systems on pseudoscorpion density, diversity and trophic niches. Soil Organisms. 93:83–96

Lindgren F, Rue H, Lindström J. 2011. An explicit link between gaussian fields and gaussian markov random fields: The stochastic partial differential equation approach. Journal of the Royal Statistical Society: Series B (Statistical Methodology). 73:423–498

Lindqvist OV. 1968. Water regulation in terrestrial isopods, with comments on their behavior in a stimulus gradient. Annales Zoologici Fennici. 5:279–311

Little C. 1983. The colonisation of land: Origins and adaptations of terrestrial animals. Cambridge University Press.

Liu M, Trugman AT, Peñuelas J, Anderegg WRL. 2024. Climate-driven disturbances amplify forest drought sensitivity. Nature Climate Change. 14:746–752

Lo N, Tong KJ, Rose HA, Ho SYW, Beninati T, Low DLT, Matsumoto T, Maekawa K. 2016. Multiple evolutionary origins of australian soil-burrowing cockroaches driven by climate change in the neogene. Proceedings of the Royal Society B: Biological Sciences. 283:20152869

Luan J, Li S, Liu S, Wang Y, Ding L, Lu H, Chen L, Zhang J, Zhou W, Han S, Zhang Y, Hättenschwiler S. 2024. Biodiversity mitigates drought effects in the decomposer system across biomes. Proceedings of the National Academy of Sciences. 121:e2313334121

Lucio-García JN, Sánchez-Reyes UJ, Horta-Vega JV, Reyes-Muñoz JL, Clark SM, Niño-Maldonado S. 2022. Seasonal and microclimatic effects on leaf beetles (Coleoptera, Chrysomelidae) in a tropical forest fragment in northeastern mexico. Zookeys. 1080:21–52

Ma H, Mo L, Crowther TW, Maynard DS, van den Hoogen J, Stocker BD, Terrer C, Zohner CM. 2021. The global distribution and environmental drivers of aboveground versus belowground plant biomass. Nature Ecology & Evolution. 5:1110–1122

Maestre FT, Delgado-Baquerizo M, Jeffries TC, Eldridge DJ, Ochoa V, Gozalo B, Quero JL, García-Gómez M, Gallardo A, Ulrich W, Bowker MA, Arredondo T, Barraza-Zepeda C, Bran D, Florentino A, Gaitán J, Gutiérrez JR, Huber-Sannwald E, Jankju M, Mau RL, Miriti M, Naseri K, Ospina A, Stavi I, Wang D, Woods NN, Yuan X, Zaady E, Singh BK. 2015. Increasing aridity reduces soil microbial diversity and abundance in global drylands. Proceedings of the National Academy of Sciences. 112:15684–15689

Maire V, Gross N, Börger L, Proulx R, Wirth C, Pontes L, Soussana J-F, Louault F. 2012. Habitat filtering and niche differentiation jointly explain species relative abundance within grassland communities along fertility and disturbance gradients. New Phytologist. 196:497–509

Malhi Y, Doughty CE, Goldsmith GR, Metcalfe DB, Girardin CAJ, Marthews TR, del Aguila-Pasquel J, Aragão LEOC, Araujo-Murakami A, Brando P, da Costa ACL, Silva-Espejo JE, Farfán Amézquita F, Galbraith DR, Quesada CA, Rocha W, Salinas-Revilla N, Silvério D, Meir P, Phillips OL. 2015. The linkages between photosynthesis, productivity, growth and biomass in lowland amazonian forests. Global Change Biology. 21:2283–2295

Manzoni S, Schimel JP, Porporato A. 2012. Responses of soil microbial communities to water stress: Results from a meta-analysis. Ecology. 93:930–938

Martin N, Glauco M, Arik H, Carl NK. 2023. Fungus and fruit consumption by harvestmen and spiders (Opiliones, Araneae): The vegetarian side of two predominantly predaceous arachnid groups. The Journal of Arachnology. 51:1–18

Martin PA, Fisher L, Pérez-Izquierdo L, Biryol C, Guenet B, Luyssaert S, Manzoni S, Menival C, Santonja M, Spake R, Axmacher JC, Yuste JC. 2024. Meta-analysis reveals that the effects of precipitation change on soil and litter fauna in forests depend on body size. Global Change Biology. 30:e17305

McCarthy JK, Mokany K, Ferrier S, Dwyer JM. 2018. Predicting community rank-abundance distributions under current and future climates. Ecography. 41:1572–1582

McKie BG, Schindler M, Gessner MO, Malmqvist B. 2009. Placing biodiversity and ecosystem functioning in context: Environmental perturbations and the effects of species richness in a stream field experiment. Oecologia. 160:757–770

McKie BG, Woodward G, Hladyz S, Nistorescu M, Preda E, Popescu C, Giller PS, Malmqvist B. 2008. Ecosystem functioning in stream assemblages from different regions: Contrasting responses to variation in detritivore richness, evenness and density. Journal of Animal Ecology. 77:495–504

Miles L, Newton AC, DeFries RS, Ravilious C, May I, Blyth S, Kapos V, Gordon JE. 2006. A global overview of the conservation status of tropical dry forests. Journal of biogeography. 33:491– 505

Millar CI, Stephenson NL. 2015. Temperate forest health in an era of emerging megadisturbance. Science. 349:823–826

Miller KB, Edgerly JS. 2008. Systematics and natural history of the australian genus *Metoligotoma* Davis (Embioptera: Australembiidae). Invertebrate Systematics. 22:329–344

Miller KB, Hayashi C, Whiting MF, Svenson GJ, Edgerly JS. 2012. The phylogeny and classification of Embioptera (insecta). Systematic Entomology. 37:550–570

Minelli A. 2011. Treatise on zoology - anatomy, taxonomy, biology. The myriapoda, volume 1. Brill.

Miranda-González R, McCune B. 2020. The weight of the crust: Biomass of crustose lichens in tropical dry forest represents more than half of foliar biomass. Biotropica. 52:1298–1308

Moore S, Adu-Bredu S, Duah-Gyamfi A, Addo-Danso SD, Ibrahim F, Mbou AT, de Grandcourt A, Valentini R, Nicolini G, Djagbletey G, Owusu-Afriyie K, Gvozdevaite A, Oliveras I, Ruiz-Jaen MC, Malhi Y. 2018. Forest biomass, productivity and carbon cycling along a rainfall gradient in west africa. Global Change Biology. 24:e496–e510

Moreno-Rueda G, Ruiz-Ruiz A, Collantes-Martín E, Arrébola JR. 2009. Relative importance of humidity and temperature on microhabitat use by land snails in arid versus humid environments. Arid Environments and wind erosion. 331–343

Moss WE, Crausbay SD, Rangwala I, Wason JW, Trauernicht C, Stevens-Rumann CS, Sala A, Rottler CM, Pederson GT, Miller BW, Magness DR, Littell JS, Frelich LE, Frazier AG, Davis KT, Coop JD, Cartwright JM, Booth RK. 2024. Drought as an emergent driver of ecological transformation in the twenty-first century. BioScience. 74:524–538

Moura RF, Tizo-Pedroso E, Del-Claro K. 2018. Colony size, habitat structure, and prey size shape the predation ecology of a social pseudoscorpion from a tropical savanna. Behavioral Ecology and Sociobiology. 72:103

Muller-Landau HC, Cushman KC, Arroyo EE, Martinez Cano I, Anderson-Teixeira KJ, Backiel B. 2021. Patterns and mechanisms of spatial variation in tropical forest productivity, woody residence time, and biomass. New Phytol. 229:3065–3087

Müller LM, Bahn M. 2022. Drought legacies and ecosystem responses to subsequent drought. Global Change Biology. 28:5086–5103

Murphy KL, Klopatek JM, Klopatek CC. 1998. The effects of litter quality and climate on decomposition along an elevational gradient. Ecological Applications. 8:1061–1071

Murphy PG, Lugo AE. 1986. Ecology of tropical dry forest. Annual Review of Ecology and Systematics. 17:67–88

Nielsen UN, Ball BA. 2015. Impacts of altered precipitation regimes on soil communities and biogeochemistry in arid and semi-arid ecosystems. Global Change Biology. 21:1407–1421

Njoroge DM, Chen S-C, Zuo J, Dossa GGO, Cornelissen JHC. 2022. Soil fauna accelerate litter mixture decomposition globally, especially in dry environments. Journal of Ecology. 110:659–672

O’Hanlon RP, Bolger T. 1999. The importance of *Arcitalitrus dorrieni* (Hunt) (Crustacea: Amphipoda: Talitridae) in coniferous litter breakdown. Applied Soil Ecology. 11:29–33

Ochoa-Hueso R, Collins SL, Delgado-Baquerizo M, Hamonts K, Pockman WT, Sinsabaugh RL, Smith MD, Knapp AK, Power SA. 2018. Drought consistently alters the composition of soil fungal and bacterial communities in grasslands from two continents. Global Change Biology. 24:2818–2827

Pan Y, Birdsey RA, Phillips OL, Jackson RB. 2013. The structure, distribution, and biomass of the world’s forests. Annual Review of Ecology, Evolution, and Systematics. 44:593–622

Parsons SA, Congdon RA. 2008. Plant litter decomposition and nutrient cycling in north queensland tropical rain-forest communities of differing successional status. Journal of Tropical Ecology. 24:317–327

Petraglia A, Cacciatori C, Chelli S, Fenu G, Calderisi G, Gargano D, Abeli T, Orsenigo S, Carbognani M. 2019. Litter decomposition: Effects of temperature driven by soil moisture and vegetation type. Plant and Soil. 435:187–200

Potapov AM, Beaulieu F, Birkhofer K, Bluhm SL, Degtyarev MI, Devetter M, Goncharov AA, Gongalsky KB, Klarner B, Korobushkin DI, Liebke DF, Maraun M, Mc Donnell RJ, Pollierer MM, Schaefer I, Shrubovych J, Semenyuk II, Sendra A, Tuma J, Tůmová M, Vassilieva AB, Chen T-W, Geisen S, Schmidt O, Tiunov AV, Scheu S. 2022. Feeding habits and multifunctional classification of soil-associated consumers from protists to vertebrates. Biological Reviews. 97:1057–1117

Powers JS, Montgomery RA, Adair EC, Brearley FQ, DeWalt SJ, Castanho CT, Chave J, Deinert E, Ganzhorn JU, Gilbert ME, González-Iturbe JA, Bunyavejchewin S, Grau HR, Harms KE, Hiremath A, Iriarte-Vivar S, Manzane E, De Oliveira AA, Poorter L, Ramanamanjato J-B, Salk C, Varela A, Weiblen GD, Lerdau MT. 2009. Decomposition in tropical forests: A pan-tropical study of the effects of litter type, litter placement and mesofaunal exclusion across a precipitation gradient. Journal of Ecology. 97:801–811

Prescott CE, Vesterdal L. 2021. Decomposition and transformations along the continuum from litter to soil organic matter in forest soils. Forest Ecology and Management. 498:119522

Ray SW, Bryan SM, Melany CF, Paul JH. 2014. Ground-dwelling beetle responses to long-term precipitation alterations in a hardwood forest. Southeastern Naturalist. 13:138–155

Read VMSJ, Hughes RN. 1987. Feeding behaviour and prey choice in *Macroperipatus torquatus* (Onychophora). Proceedings of the Royal Society of London B Biological Sciences. 230:483– 506

Reinhard J, Rowell DM. 2005. Social behaviour in an australian velvet worm, *Euperipatoides rowelli* (Onychophora: Peripatopsidae). Journal of Zoology. 267:1–7

Rentz D. 2014. A guide to the cockroaches of Australia. CSIRO publishing.

Richardson AMM, Devitt DM. 1984. The distribution of four species of terrestrial amphipods (Crustacea, Amphipoda: Talitridae) on mt. wellington, tasmania. The Australian zoologist. 21:143–156

Roberts CS, McClain EL, Seely MK, Mitchell D, Goodall VL, Henschel JR. 2025. Beetling the heat – the diurnal namib desert beetle onymacris plana cools by running. Journal of Experimental Biology. 228:jeb250379

Rue H, Martino S, Chopin N. 2009. Approximate bayesian inference for latent gaussian models by using integrated nested laplace approximations. Journal of the Royal Statistical Society: Series B (Statistical Methodology). 71:319–392

Sagi N, Hawlena D. 2024. Climate dependence of the macrofaunal effect on litter decomposition—a global meta-regression analysis. Ecology Letters. 27:e14333

Schmidt-Nielsen K, Taylor CR, Shkolnik A. 1971. Desert snails: Problems of heat, water and food. Journal of Experimental Biology. 55:385–398

Schuur EA, Matson PA. 2001. Net primary productivity and nutrient cycling across a mesic to wet precipitation gradient in hawaiian montane forest. Oecologia. 128:431–442

Schweizer M, Triebskorn R, Köhler H-R. 2019a. Snails in the sun: Strategies of terrestrial gastropods to cope with hot and dry conditions. Ecology and Evolution. 9:12940–12960

Seely MK, Louw GN. 1980. First approximation of the effects of rainfall on the ecology and energetics of a namib desert dune ecosystem. Journal of Arid Environments. 3:25–54

Seidl R, Schelhaas M-J, Lexer MJ. 2011. Unraveling the drivers of intensifying forest disturbance regimes in europe. Global Change Biology. 17:2842–2852

Shaftel R, Rinella DJ, Kwon E, Brown SC, Gates HR, Kendall S, Lank DB, Liebezeit JR, Payer DC, Rausch J, Saalfeld ST, Sandercock BK, Smith PA, Ward DH, Lanctot RB. 2021. Predictors of invertebrate biomass and rate of advancement of invertebrate phenology across eight sites in the north american arctic. Polar Biology. 44:237–257

Silva SI, MacKay WP, Whitford WG. 1985. The relative contributions of termites and microarthropods to fluff grass litter disappearance in the chihuahuan desert. Oecologia. 67:31–34

Silvertown J, Dodd ME, McConway K, Potts J, Crawley M. 1994. Rainfall, biomass variation, and community composition in the park grass experiment. Ecology. 75:2430–2437

Specht RL, Rundel PW. 1990. Sclerophylly and foliar nutrient status of mediterranean-climate plant communities in southern australia. Australian Journal of Botany. 38:459–474

Speiser B, Barker G. 2001. The biology of terrestrial molluscs. Food and feeding behaviour. 259–288

Steinberger Y, Grossman S, Dubinsky Z. 1981. Some aspects of the ecology of the desert snail *Sphincterochila prophetarum* in relation to energy and water flow. Oecologia. 50:103–108

Storch D, Bohdalková E, Okie J. 2018. The more-individuals hypothesis revisited: The role of community abundance in species richness regulation and the productivity–diversity relationship. Ecology Letters. 21:920–937

Talbot JM, Yelle DJ, Nowick J, Treseder KK. 2012. Litter decay rates are determined by lignin chemistry. Biogeochemistry. 108:279–295

Tan B, Yin R, Zhang J, Xu Z, Liu Y, He S, Zhang L, Li H, Wang L, Liu S, You C, Peng C. 2021. Temperature and moisture modulate the contribution of soil fauna to litter decomposition via different pathways. Ecosystems. 24:1142–1156

Tanaka LK, Tanaka SK. 1982. Rainfall and seasonal changes in arthropod abundance on a tropical oceanic island. Biotropica. 14:114–123

Tarli VD, Pequeno PACL, Franklin E, de Morais JW, Souza JLP, Oliveira AHC, Guilherme DR. 2014. Multiple environmental controls on cockroach assemblage structure in a tropical rain forest. Biotropica. 46:598–607

Taylor AR, Schröter D, Pflug A, Wolters V. 2004. Response of different decomposer communities to the manipulation of moisture availability: Potential effects of changing precipitation patterns. Global Change Biology. 10:1313–1324

Thiele H-U. 1977. Carabid beetles in their environments: A study on habitat selection by adaptations in physiology and behaviour. Springer-Verlag Berlin and Heidelberg GmbH & Co. K

Thomas PB, Watson PJ, Bradstock RA, Penman TD, Price OF. 2014. Modelling surface fine fuel dynamics across climate gradients in eucalypt forests of south-eastern australia. Ecography. 37:827–837

Thorpe JAT. 2024. Phylogenomics supports a single origin of terrestriality in isopods. Proceedings of the Royal Society B: Biological Sciences. 291:20241042.

Todd V. 1949. The habits and ecology of the british harvestmen (Arachnida, Opiliones), with special reference to those of the oxford district. Journal of Animal Ecology. 18:209–229

Torsekar VR, Sagi N, Daniel JA, Hawlena Y, Gavish-Regev E, Hawlena D. 2024. Contrasting responses to aridity by different-sized decomposers cause similar decomposition rates across a precipitation gradient. eLife. 13:RP93656

van Ingen LT, Campos RI, Andersen AN. 2008. Ant community structure along an extended rain forest–savanna gradient in tropical australia. Journal of Tropical Ecology. 24:445–455

Veldhuis MP, Laso FJ, Olff H, Berg MP. 2017. Termites promote resistance of decomposition to spatiotemporal variability in rainfall. Ecology. 98:467–477

Vermeij GJ, Watson-Zink VM. 2022. Terrestrialization in gastropods: Lineages, ecological constraints and comparisons with other animals. Biological Journal of the Linnean Society. 136:393–404

Vilisics F, Sólymos P, Hornung E. 2007. A preliminary study on habitat features and associated terrestrial isopod species. Contributions to Soil Zoology in Central Europe II. 195–199

Villarreal E, Martínez N, Ortiz CR. 2019. Diversity of pseudoscorpiones (Arthropoda: Arachnida) in two fragments of dry tropical forest in the colombian caribbean region. Caldasia. 41:139–151

Voigtländer K. 15 chilopoda – ecology. In: Treatise on Zoology – Anatomy, Taxonomy, Biology. The Myriapoda Volume 1. Brill. p 309–325.

Wall DH, Bradford MA, St. John MG, Trofymow JA, Behan-Pelletier V, Bignell DE, Dangerfield JM, Parton WJ, Rusek J, Voigt W, Wolters V, Gardel HZ, Ayuke FO, Bashford R, Beljakova OI, Bohlen PJ, Brauman A, Flemming S, Henschel JR, Johnson DL, Jones TH, Kovarova M, Kranabetter JM, Kutny LES, Lin K-C, Maryati M, Masse D, Pokarzhevskii A, Rahman H, SabarÁ MG, Salamon J-A, Swift MJ, Varela A, Vasconcelos HL, White DON, Zou X. 2008. Global decomposition experiment shows soil animal impacts on decomposition are climate-dependent. Global Change Biology. 14:2661–2677

Warburg MR. 1965. The evaporative water loss of three isopods from semi-arid habitats in south Australia. Crustaceana. 9:302–308

Warburg MR. 1964. The response of isopods towards temperature, humidity and light. Animal Behaviour. 12:175–186

Warburg MR, Berkovitz K. 1978. Oak-woodland pillbug *Armadillo officinalis* (Isopoda; Oniscoidea), at different humidities. Journal of Thermal Biology. 3:75–78

Warburg MR, Rankevich D, Chasanmus K. 1978. Isopod species diversity and community structure in mesic and xeric habitats of the mediterranean region. Journal of Arid Environments. 1:157– 163

Ward D, Seely MK. 1996. Competition and habitat selection in namib desert tenebrionid beetles. Evolutionary Ecology. 10:341–359

Wardle DA, Bardgett RD, Klironomos JN, Setälä H, van der Putten WH, Wall DH. 2004. Ecological linkages between aboveground and belowground biota. Science. 304:1629–1633

Weber LC, VanDerWal J, Schmidt S, McDonald WJF, Shoo LP. 2014. Patterns of rain forest plant endemism in subtropical australia relate to stable mesic refugia and species dispersal limitations. Journal of Biogeography. 41:222–238

Weeks JM. 1992. The role of coprophagy in the maintenance of body copper and zinc concentrations in talitrid amphipods (Crustacea, Amphipoda; Talitridae). Comparative Biochemistry and Physiology Part A: Physiology. 101:313–318

Wieser W. 1978. Consumer strategies of terrestrial gastropods and isopods. Oecologia. 36:191–201

Williamson DI. 1951. Studies in the biology of talitridae (Crustacea, Amphipoda): Effects of atmospheric humidity. Journal of the Marine Biological Association of the United Kingdom. 30:73–90

Zeng X, Gao H, Wang R, Majcher BM, Woon JS, Wenda C, Eggleton P, Griffiths HM, Ashton LA. 2024. Global contribution of invertebrates to forest litter decomposition. Ecology Letters. 27:e14423

Zhang-Zheng H, Adu-Bredu S, Duah-Gyamfi A, Moore S, Addo-Danso SD, Amissah L, Valentini R, Djagbletey G, Anim-Adjei K, Quansah J, Sarpong B, Owusu-Afriyie K, Gvozdevaite A, Tang M, Ruiz-Jaen MC, Ibrahim F, Girardin CAJ, Rifai S, Dahlsjö CAL, Riutta T, Deng X, Sun Y, Prentice IC, Oliveras Menor I, Malhi Y. 2024. Contrasting carbon cycle along tropical forest aridity gradients in west africa and amazonia. Nature Communications. 15:3158

Zhang D, Hui D, Luo Y, Zhou G. 2008. Rates of litter decomposition in terrestrial ecosystems: Global patterns and controlling factors. Journal of Plant Ecology. 1:85–93

Zimmer M, Kautz G, Topp W. 2005. Do woodlice and earthworms interact synergistically in leaf litter decomposition? Functional Ecology. 19:7–16

