## Supplementary material for "Moisture structures litter faunal communities through productivity and trait filtering effects"

**Supplementary Table 1.** Site coordinates, bioregion, Ecological Vegetation Class (EVC), and habitat/forest type for the 100 study sites. Forest types were defined using Ecological Vegetation Classes (EVCs) (DEECA 2026, Native Vegetation; Modelled 2005 Ecological Vegetation Classes), which classify vegetation at 1:100,000 resolution based on canopy composition, moisture availability, and plant assemblages, supplemented by a rainforest-specific layer derived from Sentinel imagery at ~10 m resolution (DEECA 2026, Rainforest Mapping for state-wide Victoria).

| **Site code** | **Latitude** | **Longitude** | **Bioregion** | **EVC** | **Habitat** |
| --- | --- | --- | --- | --- | --- |
| VIC-1-1-DRY | -37.9046285 | 145.869448 | Highlands - Southern Fall | Shrubby Foothill Forest | DRY |
| VIC-1-1-RAIN | -37.905577 | 145.811766 | Highlands - Southern Fall | Cool Temperate RF | RAIN |
| VIC-1-1-WET | -37.9142805 | 145.8351465 | Highlands - Southern Fall | Wet Forest | WET |
| VIC-1-2-DRY | -37.961141 | 146.3263605 | Highlands - Southern Fall | Shrubby Dry Forest | DRY |
| VIC-1-2-RAIN | -37.9217725 | 146.324134 | Highlands - Southern Fall | Cool Temperate RF | RAIN |
| VIC-1-2-WET | -37.9297185 | 146.334104 | Highlands - Southern Fall | Wet Forest | WET |
| VIC-1-3-DRY | -37.71748058 | 146.2767031 | Highlands - Southern Fall | Herb-rich Foothill Forest | DRY |
| VIC-1-3-MONW | -37.7524465 | 146.164829 | Victorian Alps | Montane Wet Forest | WET |
| VIC-1-3-RAIN | -37.754202 | 146.1827535 | Victorian Alps | Cool Temperate RF | RAIN |
| VIC-1-5-DAMP | -37.5371115 | 145.937434 | Highlands - Northern Fall | Damp Forest | DAMP |
| VIC-1-5-DRY | -37.5116635 | 146.001951 | Highlands - Northern Fall | Herb-rich Foothill Forest | DRY |
| VIC-1-5-WET | -37.5605675 | 145.937195 | Highlands - Southern Fall | Wet Forest | WET |
| VIC-1-6-MONW | -37.7858505 | 146.0926125 | Victorian Alps | Montane Wet Forest | WET |
| VIC-1-6-RAIN | -37.784308 | 146.088741 | Victorian Alps | Cool Temperate RF | RAIN |
| VIC-1-6-WET | -37.7860845 | 146.073861 | Highlands - Southern Fall | Wet Forest | WET |
| VIC-2-1-RAIN | -38.4473943 | 146.5385454 | Strzelecki Ranges | Cool Temperate RF | RAIN |
| VIC-2-1-WET | -38.43437466 | 146.5281024 | Strzelecki Ranges | Wet Forest | WET |
| VIC-2-2-RAIN | -38.44024025 | 146.5692358 | Strzelecki Ranges | Cool Temperate RF | RAIN |
| VIC-2-2-WET | -38.43951984 | 146.5684462 | Strzelecki Ranges | Wet Forest | WET |
| VIC-2-3-RAIN | -38.28405697 | 146.0050289 | Strzelecki Ranges | Cool Temperate RF | RAIN |
| VIC-2-3-WET | -38.28316059 | 146.0072548 | Strzelecki Ranges | Wet Forest | WET |
| VIC-2-4-RAIN | -38.53612195 | 146.3203449 | Strzelecki Ranges | Cool Temperate RF | RAIN |
| VIC-2-4-WET | -38.53460472 | 146.3215673 | Strzelecki Ranges | Wet Forest | WET |
| VIC-3-1-DAMP | -37.22225588 | 148.9400953 | Highlands - Far East | Tableland Damp Forest | DAMP |
| VIC-4-1-DRY | -37.32539126 | 148.9505497 | East Gippsland Uplands | Shrubby Dry Forest | DRY |
| VIC-4-1-RAIN | -37.30106587 | 148.9673315 | Highlands - Far East | Cool Temperate RF | RAIN |
| VIC-4-1-WET | -37.29926834 | 148.9566917 | Highlands - Far East | Wet Forest | WET |
| VIC-4-2-DAMP | -37.25275422 | 148.8464991 | Monaro Tablelands | Tableland Damp Forest | DAMP |
| VIC-4-2-RAIN | -37.30081723 | 148.8390269 | Highlands - Far East | Cool Temperate RF | RAIN |
| VIC-4-3-DAMP | -37.2542364 | 148.762491 | Highlands - Far East | Damp Forest | DAMP |
| VIC-4-3-RAIN | -37.28105402 | 148.8137629 | Highlands - Far East | Cool Temperate RF | RAIN |
| VIC-4-3-WET | -37.24273471 | 148.7946665 | Highlands - Far East | Wet Forest | WET |
| VIC-5-1-DAMP | -37.47010566 | 149.8338322 | East Gippsland Uplands | Damp Forest | DAMP |
| VIC-5-1-RAIN | -37.47272924 | 149.8376218 | East Gippsland Uplands | Warm Temperate RF | RAIN |
| VIC-5-2-DAMP | -37.43941161 | 149.7377197 | East Gippsland Uplands | Damp Forest | DAMP |
| VIC-5-2-RAIN | -37.42735954 | 149.7434947 | East Gippsland Uplands | Warm Temperate RF | RAIN |
| VIC-5-3-DAMP | -37.69405037 | 149.4489395 | East Gippsland Lowlands | Damp Forest | DAMP |
| VIC-5-3-RAIN | -37.69460308 | 149.4490582 | East Gippsland Lowlands | Warm Temperate RF | RAIN |
| VIC-5-4-RAIN | -37.52969781 | 149.6867274 | East Gippsland Lowlands | Warm Temperate RF | RAIN |
| VIC-6-1-DRY | -37.220196 | 147.8197065 | East Gippsland Uplands | Shrubby Foothill Forest | DRY |
| VIC-6-1-MONW | -37.2141915 | 147.8516655 | Victorian Alps | Montane Wet Forest | WET |
| VIC-6-1-RAIN | -37.2091485 | 147.837253 | Highlands - Southern Fall | Warm Temperate RF | RAIN |
| VIC-6-2-MOND | -37.225242 | 147.9175115 | Victorian Alps | Montane Damp Forest | DAMP |
| VIC-6-3-DRY | -37.0850175 | 147.9049215 | East Gippsland Uplands | Heathy Dry Forest | DRY |
| VIC-6-3-MOND | -37.076783 | 147.9110645 | Victorian Alps | Montane Damp Forest | DAMP |
| VIC-6-3-MONW | -37.073451 | 147.9549915 | Victorian Alps | Montane Wet Forest | WET |
| VIC-7-1-MOND | -37.217676 | 147.4352835 | Victorian Alps | Montane Damp Forest | DAMP |
| VIC-7-1-MONW | -37.222434 | 147.423234 | Victorian Alps | Montane Wet Forest | WET |
| VIC-8-1-RAIN | -37.226894 | 148.0437265 | Highlands - Southern Fall | Warm Temperate RF | RAIN |
| **Site code** | **Latitude** | **Longitude** | **Bioregion** | **EVC** | **Habitat** |
| VIC-8-1-WET | -37.1838445 | 148.0702185 | Highlands - Southern Fall | Wet Forest | WET |
| VIC-8-2-DRY | -37.339398 | 148.1024085 | East Gippsland Uplands | Shrubby Dry Forest | DRY |
| VIC-8-2-RAIN | -37.327815 | 148.0936075 | East Gippsland Uplands | Warm Temperate RF | RAIN |
| VIC-8-2-WET | -37.337533 | 148.1055605 | East Gippsland Uplands | Wet Forest | WET |
| VIC-9-1-DAMP | -37.508622 | 145.51289 | Highlands - Northern Fall | Damp Forest | DAMP |
| VIC-9-1-RAIN | -37.526849 | 145.5217165 | Highlands - Northern Fall | Cool Temperate RF | RAIN |
| VIC-9-1-WET | -37.530707 | 145.512172 | Highlands - Northern Fall | Wet Forest | WET |
| VIC-10-1-DAMP | -37.6756025 | 148.060438 | East Gippsland Lowlands | Damp Forest | DAMP |
| VIC-10-1-DRY | -37.681412 | 148.078429 | East Gippsland Uplands | Shrubby Dry Forest | DRY |
| VIC-11-1-DAMP | -37.5377905 | 148.082839 | East Gippsland Uplands | Damp Forest | DAMP |
| VIC-11-1-DRY | -37.545241 | 148.0871385 | East Gippsland Uplands | Shrubby Dry Forest | DRY |
| VIC-12-1-DAMP | -37.64677481 | 147.3964701 | East Gippsland Uplands | Shrubby Damp Forest | DAMP |
| VIC-12-1-DRY | -37.65728921 | 147.3924579 | East Gippsland Uplands | Shrubby Dry Forest | DRY |
| VIC-12-1-RAIN | -37.67702811 | 147.3908438 | East Gippsland Lowlands | Warm Temperate RF | RAIN |
| VIC-12-2-DAMP | -37.59987862 | 147.3002145 | Highlands - Southern Fall | Shrubby Damp Forest | DAMP |
| VIC-12-2-DRY | -37.61486554 | 147.3223709 | East Gippsland Uplands | Shrubby Dry Forest | DRY |
| VIC-13-1-DAMP | -37.73495 | 148.0731 | East Gippsland Lowlands | Damp Forest | DAMP |
| VIC-13-1-RAIN | -37.818 | 148.07995 | East Gippsland Lowlands | Warm Temperate RF | RAIN |
| VIC-14-1-MOND | -37.01508594 | 148.7018633 | Monaro Tablelands | Montane Damp Forest | DAMP |
| VIC-14-1-RAIN | -37.01322699 | 148.7060779 | Monaro Tablelands | Cool Temperate RF | RAIN |
| VIC-14-1-WET | -37.00565597 | 148.6615718 | Monaro Tablelands | Wet Forest | WET |
| VIC-15-1-RAIN | -37.1871432 | 148.7503775 | East Gippsland Uplands | Cool Temperate RF | RAIN |
| VIC-15-1-WET | -37.1991 | 148.76605 | Highlands - Far East | Wet Forest | WET |
| VIC-16-1-DAMP | -37.2615745 | 147.5437415 | East Gippsland Uplands | Damp Forest | DAMP |
| VIC-16-1-DRY | -37.2627765 | 147.5478015 | East Gippsland Uplands | Shrubby Dry Forest | DRY |
| VIC-16-1-MOND | -37.2777895 | 147.53068 | Victorian Alps | Montane Damp Forest | DAMP |
| VIC-16-1-WET | -37.270603 | 147.529089 | East Gippsland Uplands | Wet Forest | WET |
| VIC-17-1-DAMP | -37.13415292 | 148.8105962 | Monaro Tablelands | Tableland Damp Forest | DAMP |
| VIC-17-1-RAIN | -37.12823267 | 148.839959 | Monaro Tablelands | Cool Temperate RF | RAIN |
| VIC-17-1-WET | -37.10629167 | 148.8333892 | Monaro Tablelands | Wet Forest | WET |
| VIC-18-1-DAMP | -37.66731997 | 146.5250715 | Victorian Alps | Tableland Damp Forest | DAMP |
| VIC-18-1-RAIN | -37.67579007 | 146.52771 | Highlands - Southern Fall | Cool Temperate RF | RAIN |
| VIC-18-1-WET | -37.6802451 | 146.5390213 | Highlands - Southern Fall | Wet Forest | WET |
| VIC-19-1-DAMP | -37.53997061 | 147.002184 | Victorian Alps | Tableland Damp Forest | DAMP |
| VIC-19-1-DRY | -37.56519879 | 147.0115284 | Highlands - Southern Fall | Shrubby Dry Forest | DRY |
| VIC-19-1-WET | -37.54404387 | 147.0042456 | Highlands - Southern Fall | Wet Forest | WET |
| VIC-21-1-RAIN | -37.439388 | 147.199619 | Highlands - Southern Fall | Warm Temperate RF | RAIN |
| VIC-22-1-DRY | -37.5015025 | 147.914822 | East Gippsland Uplands | Shrubby Foothill Forest | DRY |
| VIC-22-1-RAIN | -37.504214 | 147.9205525 | East Gippsland Uplands | Warm Temperate RF | RAIN |
| VIC-22-1-WET | -37.501192 | 147.9215785 | East Gippsland Uplands | Wet Forest | WET |
| VIC-23-1-DAMP | -37.62763046 | 148.993851 | East Gippsland Lowlands | Damp Forest | DAMP |
| VIC-23-1-RAIN | -37.62458168 | 149.002003 | East Gippsland Lowlands | Warm Temperate RF | RAIN |
| VIC-24-1-DRY | -37.124005 | 147.6913995 | Highlands - Northern Fall | Heathy Dry Forest | DRY |
| VIC-24-1-MOND | -37.1265055 | 147.702884 | Highlands - Northern Fall | Montane Damp Forest | DAMP |
| VIC-24-1-MONW | -37.128261 | 147.689387 | Highlands - Northern Fall | Montane Wet Forest | WET |
| VIC-25-1-DAMP | -37.3608605 | 148.317569 | East Gippsland Uplands | Damp Forest | DAMP |
| VIC-25-1-RAIN | -37.363494 | 148.314262 | East Gippsland Uplands | Warm Temperate RF | RAIN |
| VIC-25-1-WET | -37.370954 | 148.3064115 | East Gippsland Uplands | Wet Forest | WET |
| VIC-27-1-DAMP | -37.4184495 | 149.3511217 | East Gippsland Uplands | Damp Forest | DAMP |
| VIC-27-1-DRY | -37.39418205 | 149.3435261 | East Gippsland Uplands | Shrubby Dry Forest | DRY |
| VIC-27-1-RAIN | -37.4017468 | 149.350932 | East Gippsland Uplands | Warm Temperate RF | RAIN |

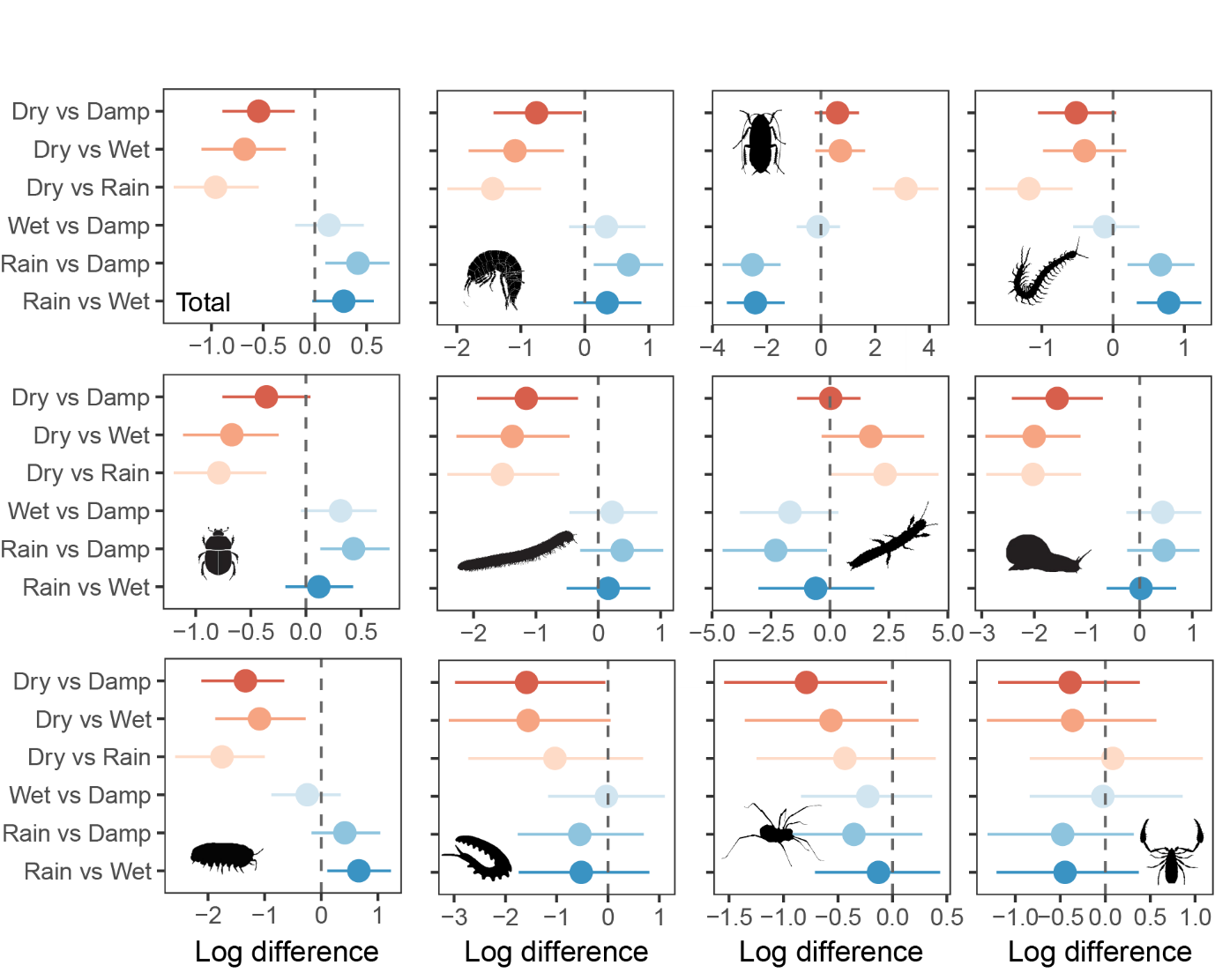

**Supplementary Figure 1.** Pairwise comparisons of the differences in the log abundance of invertebrate orders between forest habitat types, derived from 1000 posterior samples of the fitted INLA model. Panels show the posterior mean (point) and 95% credible interval (horizontal line) for each contrast for Total invertebrates, Amphipoda, Blattodea, Chilopoda, Coleoptera, Diplopoda, Embioptera, Gastropoda, Isopoda, Onychophora, Opiliones, and Pseudoscorpiones respectively. The point and errors are posterior means and credible intervals representing the relative difference between the first and second named habitats. The dashed vertical line at zero indicates no difference between habitats; contrasts whose credible interval does not overlap zero provide evidence of a meaningful difference in abundance. All differences are expressed on the natural log scale, such that exponentiating a contrast gives the ratio of expected abundances between the two habitats.

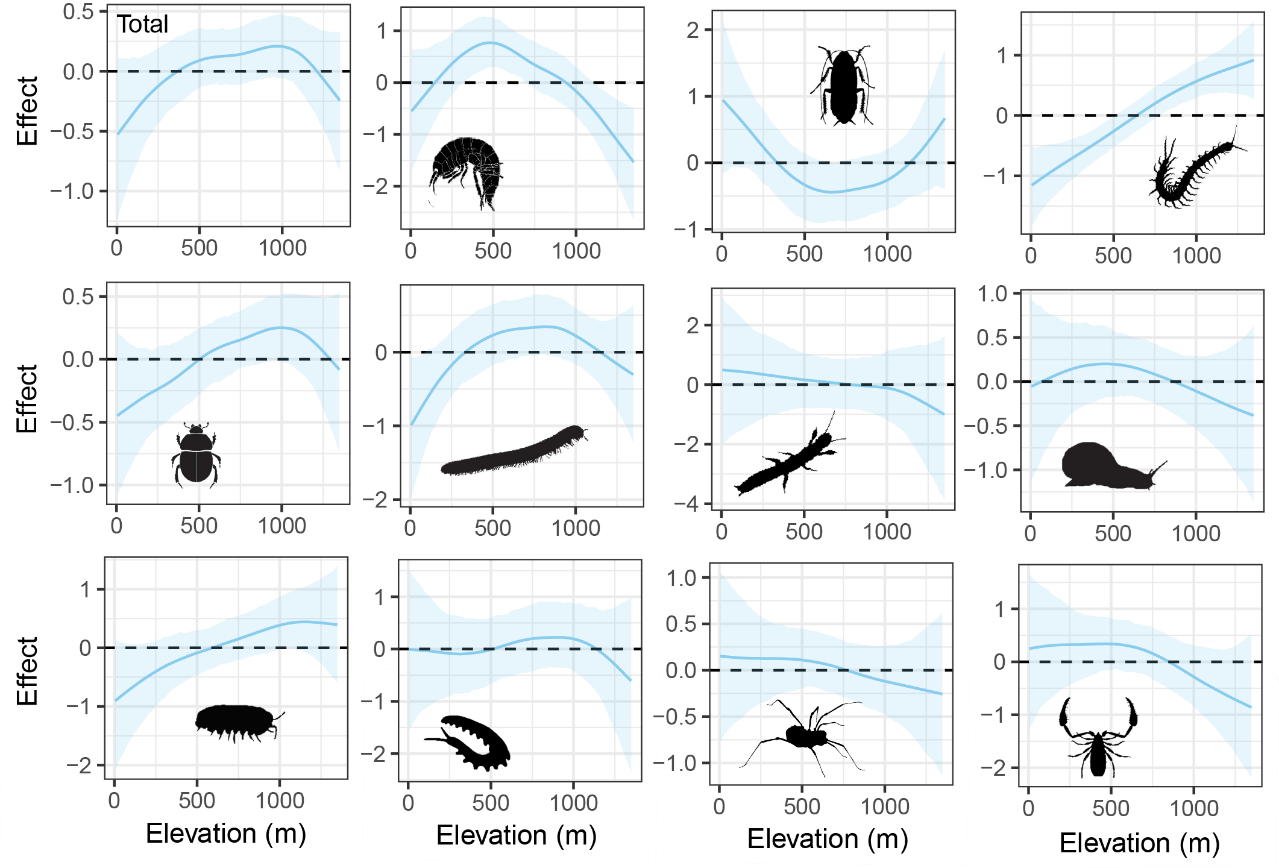
**Supplementary Figure 2.** Estimated effect of elevation (m) on invertebrate abundance for total invertebrates and each of the 11 taxonomic orders, from the Bayesian INLA model. Solid lines show the model's estimated relationship between elevation and abundance; shaded ribbons show the 95% credible interval (uncertainty range) around that estimate. Values above zero indicate higher-than-average abundance at that elevation value, and values below zero indicate lower-than-average abundance (log scale).

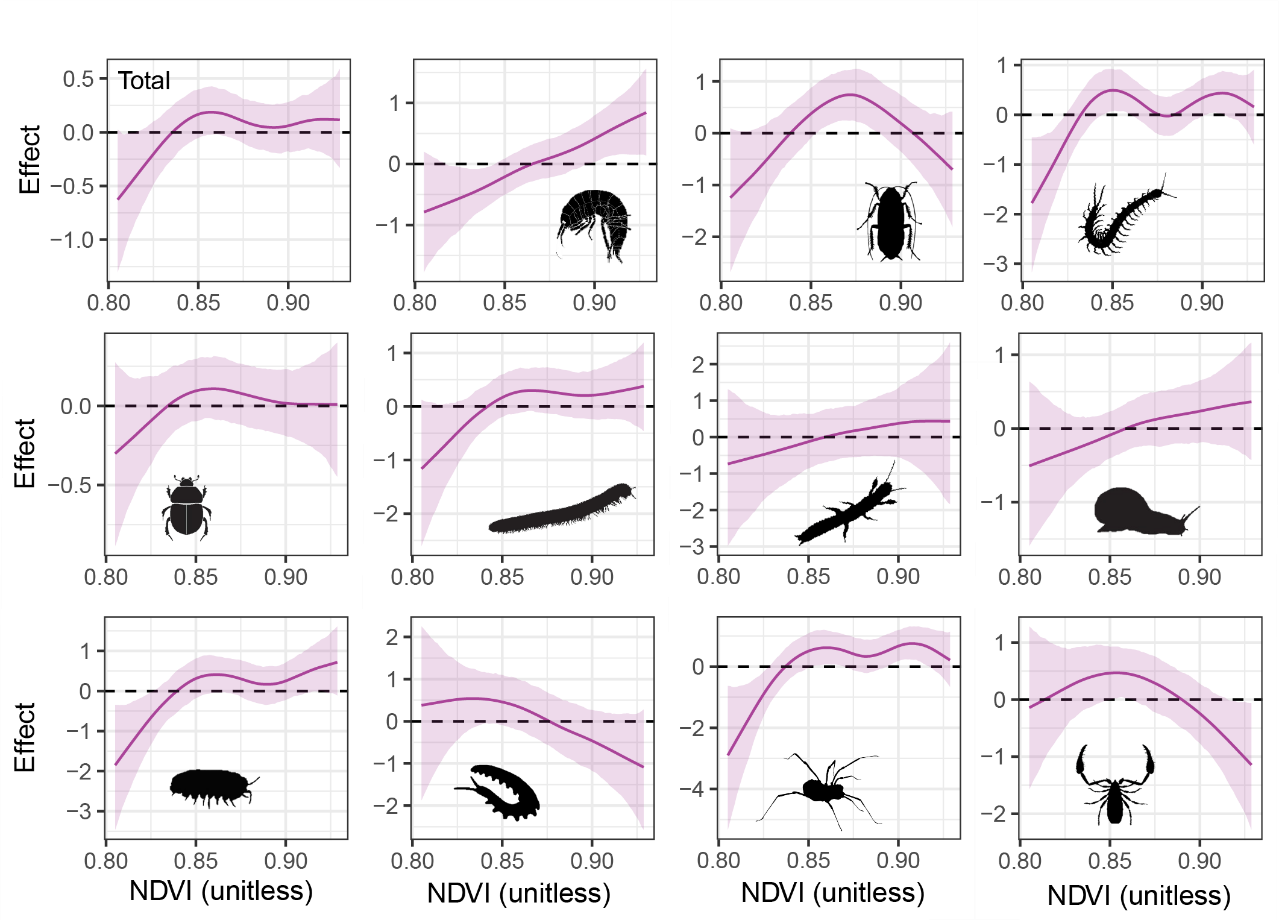

**Supplementary Figure 3.** Estimated effect of NDVI (Normalized Difference Vegetation Index; a unitless proxy for vegetation greenness/cover) on invertebrate abundance for total invertebrates and each of the 11 taxonomic orders, from the Bayesian INLA model. Solid lines show the model's estimated relationship between NDVI and abundance; shaded ribbons show the 95% credible interval (uncertainty range) around that estimate. Values above zero indicate higher-than-average abundance at that NDVI value, and values below zero indicate lower-than-average abundance (log scale).
